# Glypicans specifically regulate Hedgehog signaling through their interaction with Ihog in cytonemes

**DOI:** 10.1101/2020.11.04.367797

**Authors:** Eléanor Simon, Carlos Jiménez-Jiménez, Irene Seijo-Barandiarán, Gustavo Aguilar, David Sánchez-Hernández, Adrián Aguirre-Tamaral, Laura González-Méndez, Pedro Ripoll, Isabel Guerrero

## Abstract

The conserved family of Hedgehog (Hh) signaling proteins plays a key role in cell-cell communication in development, tissue repair and cancer progression. These proteins can act as morphogens, inducing responses dependent on the ligand concentration in target cells located at a distance. Hh proteins are lipid modified and thereby have high affinity for membranes, which hinders the understanding of their spreading across tissues. Direct contact between cell membranes by filopodia-like structures (also known as cytonemes) could be the simplest explanation for Hh dispersal. To better understand this signaling mechanism, we have analyzed in *Drosophila* the interaction between the glypicans that, besides for other pathways, are necessary for Hh signaling, plus the adhesion molecules and Hh coreceptors Ihog and Boi. We describe that glypicans (Dally and Dally-like protein) are required to maintain Ihog, but not Boi, protein levels. We also show that ectopic Ihog stabilizes cytonemes through its interaction with glypicans, and we determine that two Ihog fibronectin III domains are essential for this interaction. Our data suggest that this interaction with Ihog in cytonemes confers the specificity of glypicans for Hh signaling.

## Introduction

The Hedgehog (Hh) signaling pathway has a conserved central role in cell-cell communication during tissue patterning, stem cell maintenance and cancer progression (Briscoe and Thérond, 2013). During development, Hh ligand is usually released from a localized source inducing concentration-dependent cellular differentiation and/or proliferation responses in the surrounding cells. Hh is synthetized as a precursor molecule that undergoes two post-translational lipid modifications: by cholesterol (Porter, Young and Beachy, 1996) and by palmitic acid (Pepinsky *et al*., 1998), which associates it to cell membranes restricting its free dispersion through the extracellular milieu (Peters *et al*., 2004). Several mechanisms have been proposed to explain Hh dispersion (Lewis *et al*., 2001); among these, the transport by filopodia-like structures (also known as cytonemes) is an attractive model, since they facilitate direct contact between cell membranes.

Cytonemes have been described as dynamic actin-based protrusive structures that deliver or uptake many signaling proteins in different biological contexts (Ramírez-Weber and Kornberg, 1999); reviewed in (González-Méndez, Gradilla and Guerrero, 2019). In *Drosophila*, Hh has been shown to be localized along cytonemes in the germline stem cells niche (Rojas-Ríos, Guerrero and González-Reyes, 2012), in abdominal histoblast nests and in wing imaginal discs (Callejo *et al*., 2011; Bilioni *et al*., 2013; Gradilla *et al*., 2014; Chen *et al*., 2017), and it have been described a spatial and temporal correlations between Hh gradient establishment and cytoneme formation (Bischoff *et al*., 2013; González-Méndez, Seijo-Barandiarán and Guerrero, 2017). As the *Drosophila* Hh (Gradilla *et al*., 2014), the vertebrate homolog Sonic hh, has also been visualized *in vivo* in vesicle-like structures moving along filopodia of Shh-producing cells in the chick limb bud (Sander *et al*., 2013). Although the importance of cytonemes in the coordination of growth and patterning is well documented (Ali-Murthy and Kornberg, 2016; González-Méndez, Gradilla and Guerrero, 2019; Zhang and Scholpp, 2019), much less is known about the intrinsic and extrinsic cues regulating their establishment and dynamics.

In larval wing imaginal discs and pupal abdominal histoblast nests, Hh is produced and secreted by the Posterior (P) compartment cells and transported towards the receiving Anterior (A) compartment cells, resulting in a graded Hh distribution in the A compartment. Cytonemes emanating at the basal surface of the polarized epithelia guide Hh delivery directly from P to A cytonemes at contact sites where Hh reception takes place (Bischoff *et al*., 2013; Chen *et al*., 2017; González-Méndez, Seijo-Barandiarán and Guerrero, 2017; González-Méndez *et al*., 2020). To activate its targets in a concentration-dependent manner, Hh binds to its receptor complex formed by the canonical receptor Patched (Ptc), the co-receptors Interference hedgehog (Ihog) and Brother of Ihog (Boi) (Yao *et al*., 2006), and the membrane-anchored glypicans Dally-like protein (Dlp) and Dally (Desbordes and Sanson, 2003; Lum, Yao, *et al*., 2003; C. H. Han *et al*., 2004; Williams *et al*., 2010). All these proteins have been described associated to Hh presenting (Bilioni *et al*., 2013; Bischoff *et al*., 2013) and receiving cytonemes (Chen *et al*., 2017; González-Méndez, Seijo-Barandiarán and Guerrero, 2017).

The glypicans Dlp and Dally have been described as being required for cytoneme elongation and/or attachment in the wing imaginal discs (Huang and Kornberg, 2016; González-Méndez, Seijo-Barandiarán and Guerrero, 2017). Glypicans are composed of polysaccharide chains of heparin sulfate attached to a core protein bound to the membrane via a GPI anchor and they have a generalized, but not uniform, expression in the wing imaginal disc. Glypicans regulate several morphogenetic gradients (ie. Hh, Wingless, decapentaplegic, FGF, JAK/STAT signaling) (Yan *et al*., 2009; Nakato and Li, 2016)); however, how glypicans achieve their specifity for each signal is not known. Both Dlp and Dally are needed for Hh reception and to maintain Hh extracellular levels. Indeed, mutant clones for the glypican Dlp fail to transduce Hh (Desbordes and Sanson, 2003; Lum, Zhang, *et al*., 2003; C. H. Han *et al*., 2004; Williams *et al*., 2010) and impair cytoneme formation in the air sac primordium (Huang and Kornberg, 2016).

To date, the co-receptors Ihog and Boi are known to have redundant functions because Hh signaling is only blocked when both are simultaneously absent in the Hh receiving cells (Zheng *et al*., 2010). Ihog and Boi are also independently needed for the maintenance of normal Hh levels in Hh-producing cells (Yan *et al*., 2010). Both proteins are expressed ubiquitously in the wing disc, although their protein levels are reduced at the A/P boundary in response to Hh signaling. Ihog and Boi share sequence similarities, they are Type 1 transmembrane proteins with four Ig and two Fibronectine type III (FNIII) extracellular domains, and an undefined intracellular domain (Yao *et al*., 2006). Ihog is preferably directed towards basal membranes (Zheng *et al*., 2010; Bilioni *et al*., 2013; Hsia *et al*., 2017) and it has the ability to stabilize cytonemes when over-expressed, clearly influencing cytoneme dynamics (Bischoff *et al*., 2013; González-Méndez, Seijo-Barandiarán and Guerrero, 2017).

In this work, we further analyze the interaction of glypicans with Ihog and Boi and the role of these interactions on cytoneme dynamics and Hh signaling. We describe that whereas Ihog and Boi functions are not needed for the maintenance of glypicans levels, glypicans are required to maintain Ihog protein levels. In addition, we observe that ectopic Ihog, but not Boi, stabilizes cytonemes. We further dissected the functional domains of Ihog responsible for the interaction with glypicans, Hh and Ptc, as well as for cytoneme stability and Hh gradient formation. We describe that the Hh-binding Fn1 domain of Ihog (McLellan *et al*., 2006) is crucial for glypican interaction and cytoneme stabilization, though the amino-acids involved are different from those responsible for the Ihog/Hh interaction. Similarly, the Ptc-interacting Fn2 domain (Zheng *et al*., 2010) is also key for Ihog interaction with glypicans as well as for the stabilization of cytonemes. We conclude that Ihog interaction with glypicans regulates cytoneme behavior through both FNIII domains, being these interactions key for Hh gradient formation while providing the specificity of glypican function for Hh signaling.

## Results

### Glypicans interact with Ihog to stabilize it at the plasma membranes

Previous works have already described a genetic interaction between Ihog, Boi and Dlp (Yao *et al*., 2006) as well as an interaction between glypicans and Ihog, revealed by the ectopic Ihog recruitment of glypicans in wing disc cells (Bilioni *et al*., 2013). Here we have further investigated such interaction. We found that Dlp and Dally do not change their protein levels in *boi^-/-^ ihog^-/-^* double mutant clones (Fig. 1A,B). Conversely, in double *ttv^-/-^ botv^-/-^* mutant clones, responsible of synthesizing the Heparan Sulfate chains of the glypicans Dally and Dlp, Ihog protein is barely present at the plasma membrane of the disc epithelium (Fig. 1C). In addition, we observed that this downregulation is not significant within *dlp* or *dally* single mutant clones (Fig. 1E,F) but it is again present in *dally dlp* double mutant clones (Fig. 1D). Thus, this effect on Ihog requires of both Dally and Dlp proteins; it seems to be dose-dependent since in sister clones (homozygous *dlp^+/+^dally*^+/+^ wild type cells contiguous to homozygous mutant clones *dlp^-/-^ dally^-/-^*) there are higher levels of Ihog than in the heterozygous *dlp^-/+^ dally^-/+^* background cells (Fig. 1 C,D arrowheads), further indicating that the modulation of Ihog levels might occur through protein stabilization at the plasma membranes.

**Figure 1.**
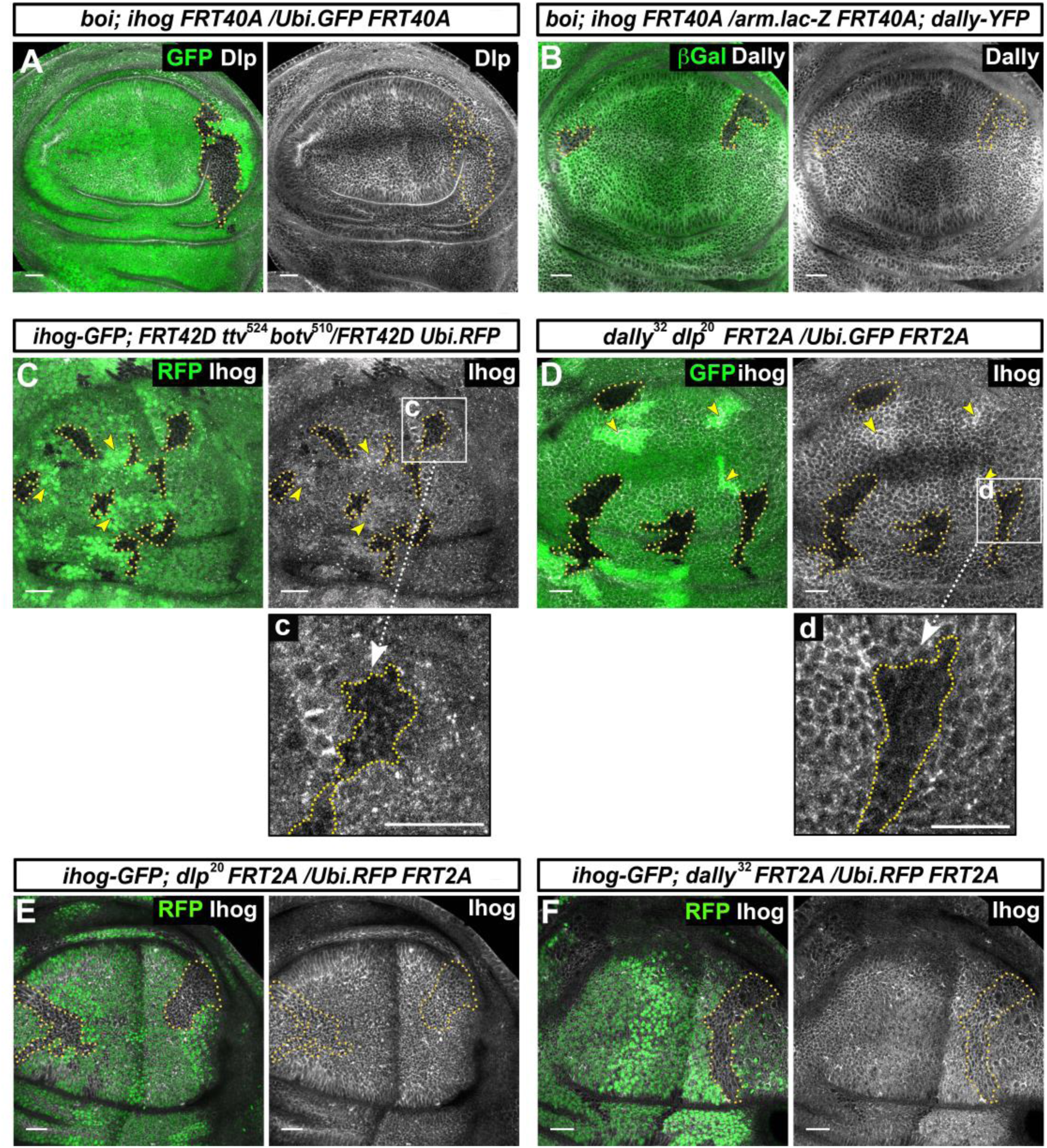
Glypicans regulate Ihog presence at the plasma membrane. A, B) *boi*; *ihog^Z23^* double mutant clones (labelled by the lack of GFP) do not affect the expression of the glypicans Dlp (A) and Dally (B) (grey channel). C) Ihog levels (BacIhog GFP, grey channel) decrease in *ttv^524^ botv^510^* double mutant clones (green channel); c) enlargement of one of the clones. D) Ihog levels (labelled with anti Ihog antibody, grey channel) decrease in *dally^32^ dlp^20^* double mutant clones (green channel); d) enlargement of one clone. Note that the homozygous wild type sister clones (more intense green) positively modulate Ihog levels (arrowheads in C and D). E,F) *dlp^20^* (E) and *dally^32^* (F) single mutant clones (green channel) do not affect Ihog levels (anti Ihog antibody, grey channel, arrows). Scale bar: 20 μm.

Interestingly, in the case of Boi there is no evidence for a similar modulation. In contrast with the strong downregulation of Ihog, silencing of Dally and Dlp through RNAi expression provokes a very slight upregulation of Boi (Supplementary Fig. 1B,B’). In the same direction, we observed an increase of Boi levels when Ihog is downregulated by RNAi expression (Fig. 1C,C’), which indicates that the slight upregulation observed on Boi in *dlp dally* mutants might not be directly mediated by glypicans. In summary, these data suggest a novel functional interaction of Ihog, but not of Boi, with Dally and Dlp to maintain Ihog protein levels at the plasma membrane.

### Ihog functional domains implicated in the Ihog-glypican interaction

We have investigated the function of the various Ihog protein domains in Ihog-Dally and Ihog-Dlp interactions. To this end, we generated transgenic lines to ectopically express Ihog proteins carrying deletions for the different extracellular domains or containing only the intracellular C-terminal fragment, each fused to the red fluorescent protein (RFP) or to hemagglutinin (HA) epitope. The following constructs were generated: Ihog lacking the four Ig domains (*UAS.ihogΔIg-RFP*), the two FNIII domains *(UAS.ihogΔFn1,2-RFP*), the Fn1 domain (*UAS.ihogΔFn1-RFP*), the Fn2 domain (*UAS.ihogΔFn2-HA)* and Ihog containing only the intracellular Ihog C-terminus (*UAS.ihogCT-RFP*), including in both constructs the transmembrane domain. The molecular weights corresponding to the generated Ihog mutant forms are shown by western blot (Supplementary Fig 2).

**Figure 2.**
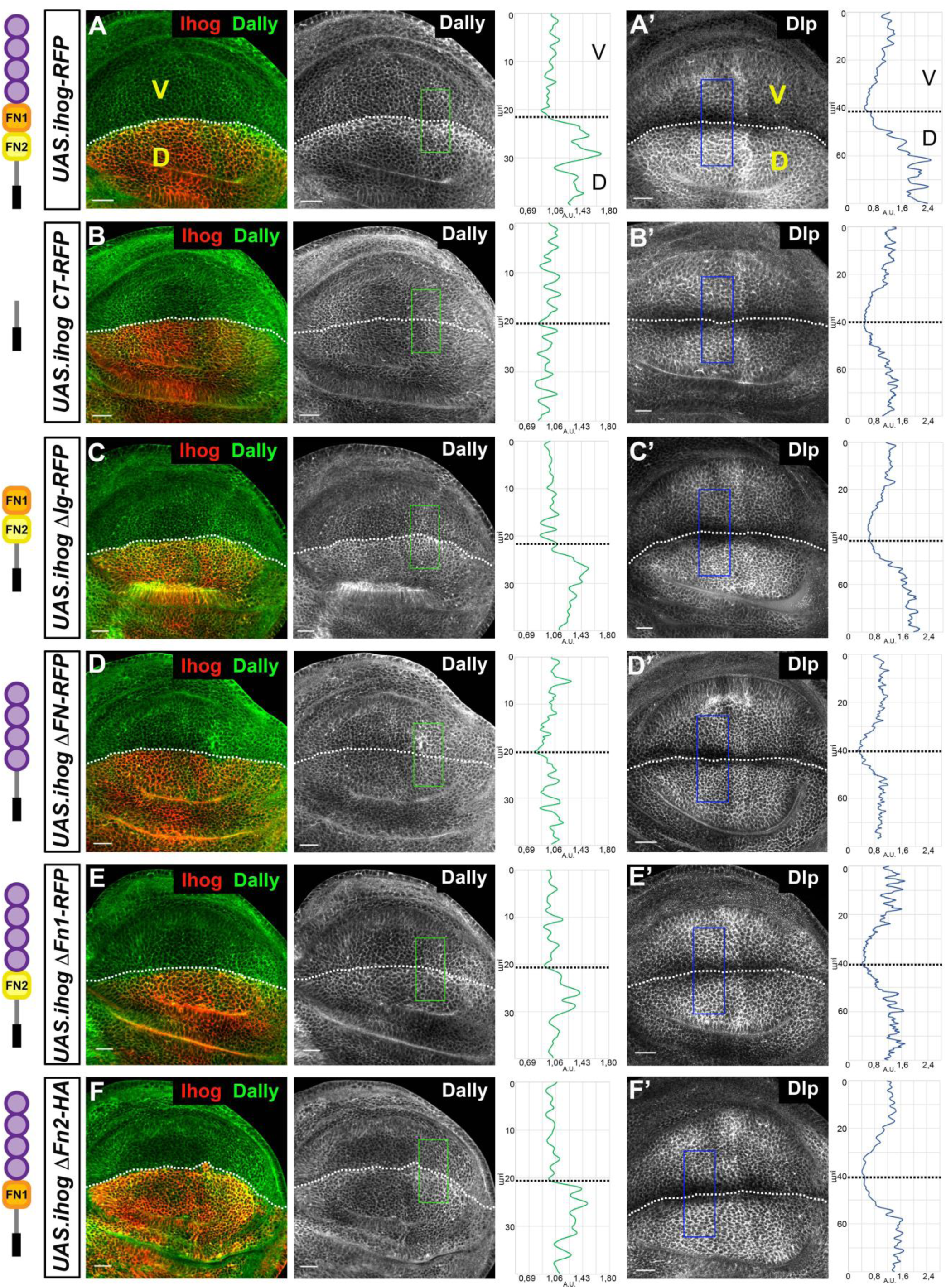
Role of Ihog domains in the interaction with glypicans. A-F) Glypicans accumulation in: *apGal4 tubGal80^ts^/+; UAS.ihog-RFP/+* (A-A’); *apGal4 tubGal80^ts^/ UAS.ihogCT-* RFP/+ (B-B’); *apGal4 tubGal80^ts^/ +: UAS.ihogΔIg-RFP/ +* (C-C’); *apGal4 tubGal80^ts^/+; UAS.ihogΔFn-RFP/ +* (D-D’); *apGal4 tubGal80^ts^/ +; UAS.ihogΔFn1-RFP/+* (E-E’); *apGal4tubGal80^ts^/+; UAS.ihogΔFn2-HA/+* (F-F’) wing discs after 30 hours at the restricted temperature. Each image incorporates at its side a plot profile (taken from the framed area in each image), indicating the relative intensity fluorescence for each glypican. Note that IhogCT-RFP and IhogΔFN-RFP do not build-up either Dally (B, D) or Dlp (B’, D’). However, in ectopic expression of a partial deletions of the FN-type III domains (IhogΔFn1-RFP and IhogΔFn2-HA) an accumulation of Dally (E, F), but not of Dlp (E’, F’), is observed although in lower levels than those of the ectopic Ihog-RFP (A, A’). Scale bar: 20 μm. Discs are oriented with dorsal part down (D), ventral up (V) and posterior right. Plot profiles were done over a ROI of size of 40μm X 20μm for Dally and 80 μm X 20 μm for Dlp.

The tests for the protein domain(s) responsible for the Ihog/glypican interaction were done through analysis of the glypicans Dally and Dlp build-up effect after overexpressing the Ihog mutant variants in the dorsal compartment of the wing imaginal disc. First, we found no accretion of Dally or Dlp after ectopic expression of *UAS.ihogCT-RFP* (Fig. 2B,B’). This result excludes the intracellular domain of Ihog from this interacting function. The ectopic expression of *UAS.ihogΔIg-RFP* in the dorsal part of the wing disc accumulates glypicans at levels similar to the expression of the full-length Ihog form (Fig. 2C,C’), while the ectopic expression of *UAS.ihogΔFN-RFP* does not result in Dally or Dlp build-up (Fig. 2D,D’), confirming the implication of the extracellular fragment and specifically identifying the fibronectin domains as responsible for the glypican interaction. We then determined which of the two FNIII domains was responsible for the Ihog-glypican interaction. Expression of *UAS.ihogΔFn1-RFP* or *UAS.ihogΔFn2-HA* results in low levels of Dally accretion and no Dlp build-up (Fig. 2E,E’ and F,F’) in comparison with the expression of wild type *UAS.IhogRFP* (Fig. 2A,A’), indicating that both FNIII domains are required for glypicans-Ihog interactions.

### The extracellular Ihog Fn2 domain interacts with both Hh and glypicans through different amino-acid sequences

Both Ihog and Boi have a demonstrated role in the maintenance of extracellular Hh levels in the Hh producing cells since the extracellular levels of Hh decrease in wing discs after the loss of function of Ihog, Boi or both. Furthermore, Hh builds-up at the plasma membrane in fixed wing imaginal discs overexpressing Ihog (Fig. 3A) (Yan *et al*., 2010; Callejo *et al*., 2011; Bilioni *et al*., 2013). The Ihog-Hh interaction was also shown to take place through the Ihog Fn1 domain (McLellan *et al*., 2006; Yao *et al*., 2006) and, as expected, we found that the expression of either *UAS-ihogΔFN-RFP* or *UAS-ihogΔFn1-RFP* does not result in Hh accumulation (Fig. 3F and Fig. 3B, respectively).

**Figure 3.**
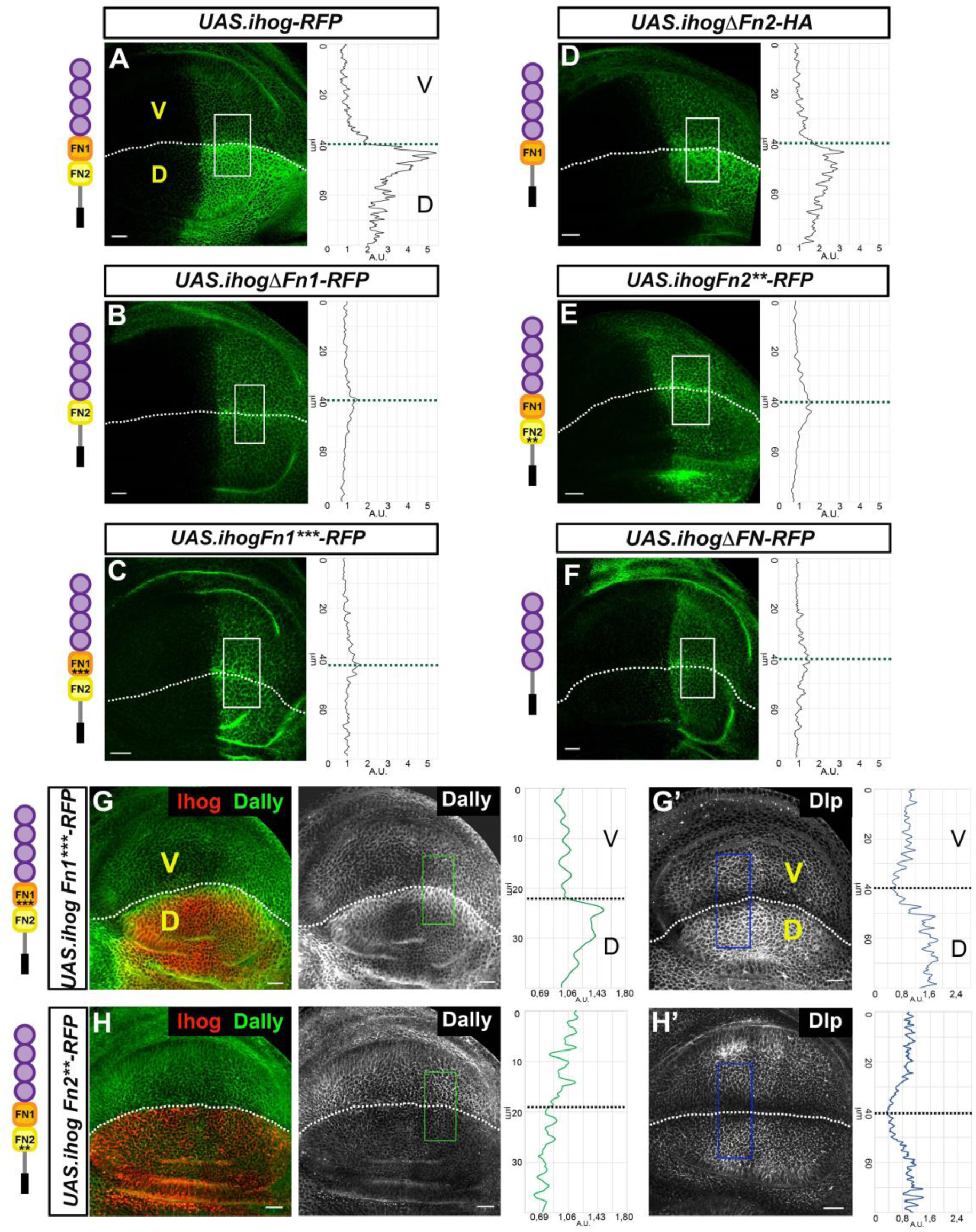
Effects of Ihog Fn1 and Fn2 deletion and point mutations on Hh retention and glypican interaction. A-F) Hh (BacHhGFP) expression in wing discs after 30 hours at the restricted temperature in *apGal4 BacHhGFP/+; UAS.ihog-RFP/tubGal80^ts^* (A); *apGal4 BacHhGFP/+; UAS.ihogΔFn1-RFP/tubGal80^ts^* (B); *apGal4 BacHhGFP/+; UAS.ihogFn1***-RFP/tubGal80^ts^* (C); *apGal4 BacHhGFP/+; UAS.ihogΔFn2-HA/tubGal80^ts^* (D); *apGal4 BacHhGFP/+; UAS.ihogFn2**-RFP/tubGal80^ts^* (E) and *apGal4 BacHhGFP/+; UAS.ihogΔFN-RFP/tubGal80^ts^* (F). G-H) Accumulation of glypicans after 30 hours at the restricted temperature in *apGal4 tubGal80^ts^/ +; UAS.ihogFn1***-RFP/+* (G, G’) and in *apGal4 tubGal80^ts^/ +; UAS.ihogFn2**-RFP/+* wing discs (H, H’). Images incorporate at their side a plot profile (taken from the framed area in each image), indicating a modulation of each glypican levels. Plot profiles where done over ROI of size 80μm X 30μm for Hh, 40μm X 20μm for Dally and 80μm X 20μm for Dlp. Note that Ihog Fn1*** does not interact with Hh (C) and builds-up Dally (G) and Dlp (G’). By contrast, Ihog Fn2** does not accumulate Dlp and but decreases the levels of Dally (H, H’). Scale bar: 20 μm. Discs are oriented with dorsal part down (D), ventral up (V) and posterior right.

To test the sufficiency of Ihog Fn1 domain for the interaction with Hh, we targeted it to the membrane by fusing it to the extracellular domain of CD8 (*UAS.CD8.Fn1-Cherry*). Expression of this construct in the dorsal compartment resulted in a very small increase of Hh levels (Supplementary Fig. 3C). This result, together with the fact that deletion of neither Fn1 nor Fn2 accumulates Dlp and slightly increase Dally levels (Fig. 3E,E’ and F,F’), suggests the need for Ihog-glypicans interaction in the Ihog Fn1-Hh recruitment.

To further analyze the potential Fn1 domain sequences responsible for Hh and/or glypican accumulation, we made an *in silico* prediction of the amino-acid residues responsible for the Ihog-Hh interaction based on structural analysis of Ihog (McLellan *et al*., 2008) and generated an Ihog variant carrying point mutation substitutions in three residues (D558N, N559S, E561Q) previously described to interact with Hh via hydrogen bonds (McLellan *et al*., 2006). As anticipated, overexpression of this Ihog form (*UAS.IhogFn1***-RFP*) does not accumulate Hh, probably as a result of the loss of the Ihog-Hh interaction due to at least one of the three mutated amino-acids (Fig. 3C). We next analyzed the interaction of the same Ihog mutant form, *UAS.IhogFn1\*\*\**, with glypicans and observed that both Dally and Dlp build-up (Fig. 3G, G’) at a level similar to that following overexpression of the wild type Ihog (Fig. 2A, A’), indicating that the Fn1 amino-acids that interact with glypicans are specific and different from those that carry out the Ihog-Hh interaction.

We observed no difference in Hh build-up after expression of either the wild type Ihog or the form lacking the Ig domains (*UAS.ihogΔIg-RFP)* (Supplementary Fig. 3A), while the expression of *UAS-ihog.ΔFn2-HA* still builds-up Hh (Fig. 3D) although at noticeably lower levels than the expression of the wild type Ihog (Fig 3A). This last reduction could result from a diminished interaction of Ihog.ΔFn2-HA with glypicans (Fig. 2F), as Dally is needed for Hh stability at the plasma membrane (Bilioni *et al*., 2013). Finally, as expected, Hh accretion is not observed when expressing only the intracellular part of the protein (Supplementary Fig. 3B). Hence, the Ihog interaction with Hh involves the Fn1 and, maybe partially, the Fn2 domains but not the Ig and C-terminal domains.

### Ihog FnIII domains influence Hh reception and gradient formation

Previous research had described the role of the Fn2 domain of Ihog as enhancing Hh protein binding in cells co-expressing its receptor Ptc (Yao, Lum and Beachy, 2006) through Ptc interaction with two specific Fn2 amino-acid residues (Zheng *et al*., 2010). Since Dlp and, probably, Dally are also part of the Hh reception complex, we next studied if these amino-acids could also be implicated in the interaction of Fn2 with Hh and the two glypicans. We generated an Ihog variant carrying point mutations in the two amino-acid residues (K653 and Q655) (Zheng *et al*., 2010) and fused it to the red fluorescent protein (*UAS.ihogFn2**-RFP*). Interestingly, the expression of this Ihog mutant form does not accumulate Hh (Fig. 3E), Dally or Dlp (Fig. 3H, H’). To our surprise, this is different from the expression of *UAS.ihogΔFn2* (Fig. 3D), deficient for the entire Fn2 domain, which still recruits Hh. Moreover, the expression of *UAS.ihogFn2\*\** with or without RFP decreases the amount of endogenous Dally (Fig. 3G and Supplementary Fig. 4A respectively), probably acting as dominant negative in the recruitment of Dally and thus indirectly affecting Hh stabilization.

To further investigate the effect of *UAS.ihogΔFn2-HA* and *UAS.ihogFn2**-RFP* variants on Hh reception, we overexpressed the wild type Ihog and these two mutant forms in the Ptc expressing cells but only in the ventral side of the disc. To this end we used *LexAop Gal80; apLexA Ptc Gal4* driver, thus maintaining the dorsal side as internal wild type control (Fig. 4A), and analyze the expression of the high threshold target Ptc, and the low threshold target Cubitus interruptus (Ci). After ectopic Ihog RFP expression, Hh reception is slightly affected, resulting in a flattened gradient responses by extending Ptc expression while reducing its highest levels; and also observing an extension of Ci expression (Fig. 4B; Supplementary Fig 5A, see also (Yan *et al*., 2010)). The flattening effect is clearer when expressing *UAS.ihogΔFn2-HA* (Fig. 4C; Supplementary Fig. 5B), indicating that the presentation of Ptc to the membrane is probably disturbed, also inferred by the extension of Ci expression. However, no effect is detected after the expression of *UAS-ihogFn2**-RFP,* that carries the two Fn2 point mutations (Fig. 4D, Supplementary Fig. 5C), although it had been proposed that its interaction with Ptc was reduced (Zheng *et al*., 2010).

**Figure 4.**
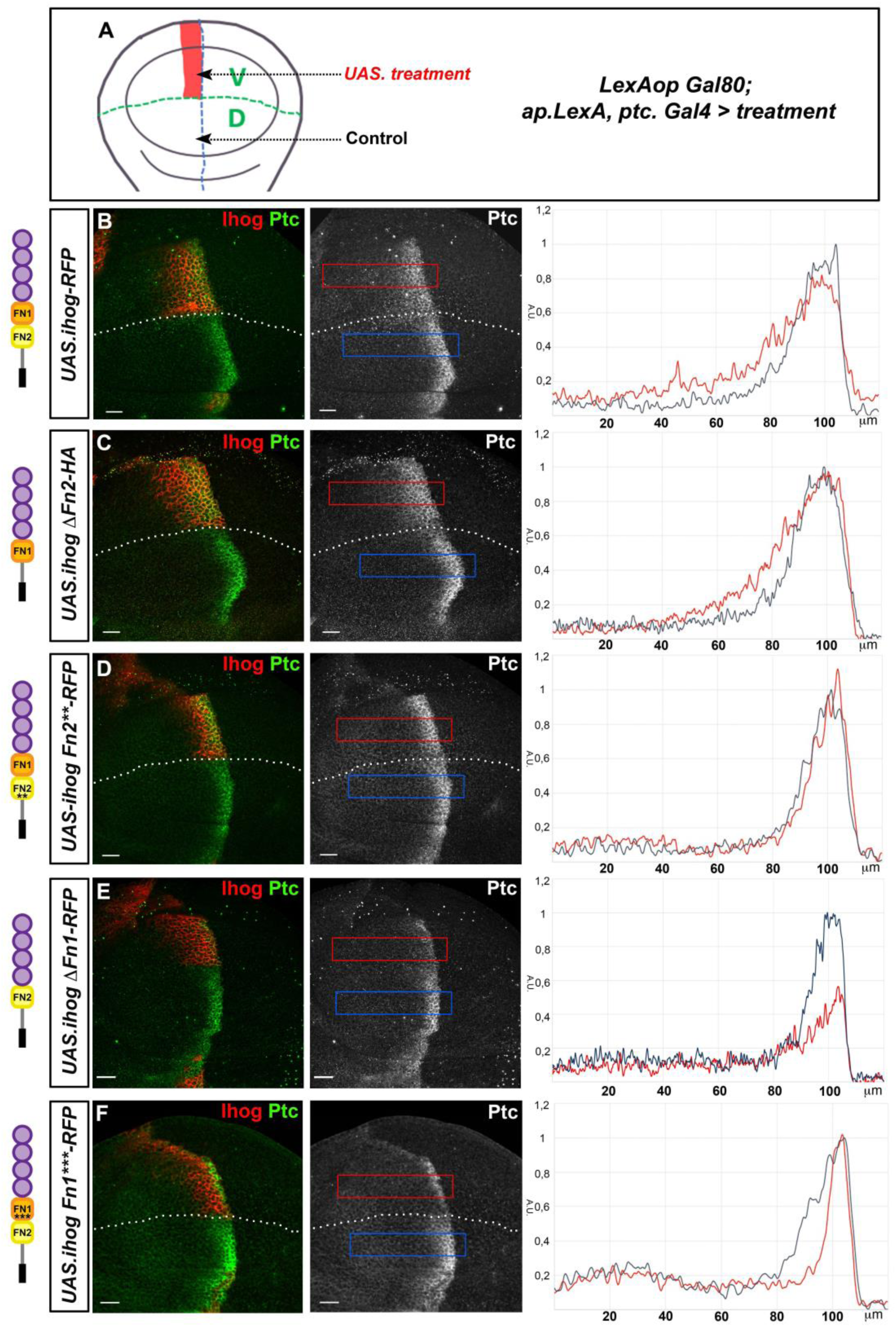
Role of the Ihog domains on Hh gradient formation. A) Scheme depicting the *LexAopGal80; apLexA ptcGal4* expression domain in the ventral part of the wing disc. B-F) Ptc expression in wing discs after induction for 28 hours of the transgenes: *UAS.ihog-RFP* (B); *UAS.ihogΔFn2-HA* (C); *UAS.ihogFn2**RFP* (D); *UAS.ihogΔFn1-RFP* (E); and *UAS.ihogFn1***RFP* (F) using the multiple driver *LexAopGal80; apLexA ptcGal4/+; tubGal80^ts^/+.* Each image incorporates plots of the fluorescence intensity of Ptc expression (red) in the ventral side of the wing disc. The dorsal side is used as internal control for Ptc wild type expression (blue). Note that Ihog *Δ*Fn1 abrogates the Hh reception while Ihog Fn1*** keeps signaling in an anterior cell row near the A/P border. Scale bar: 20 μm. Discs are oriented with dorsal part down and posterior right. Plot profiles were performed over a ROI of size 120μm X 20μm.

**Figure 5.**
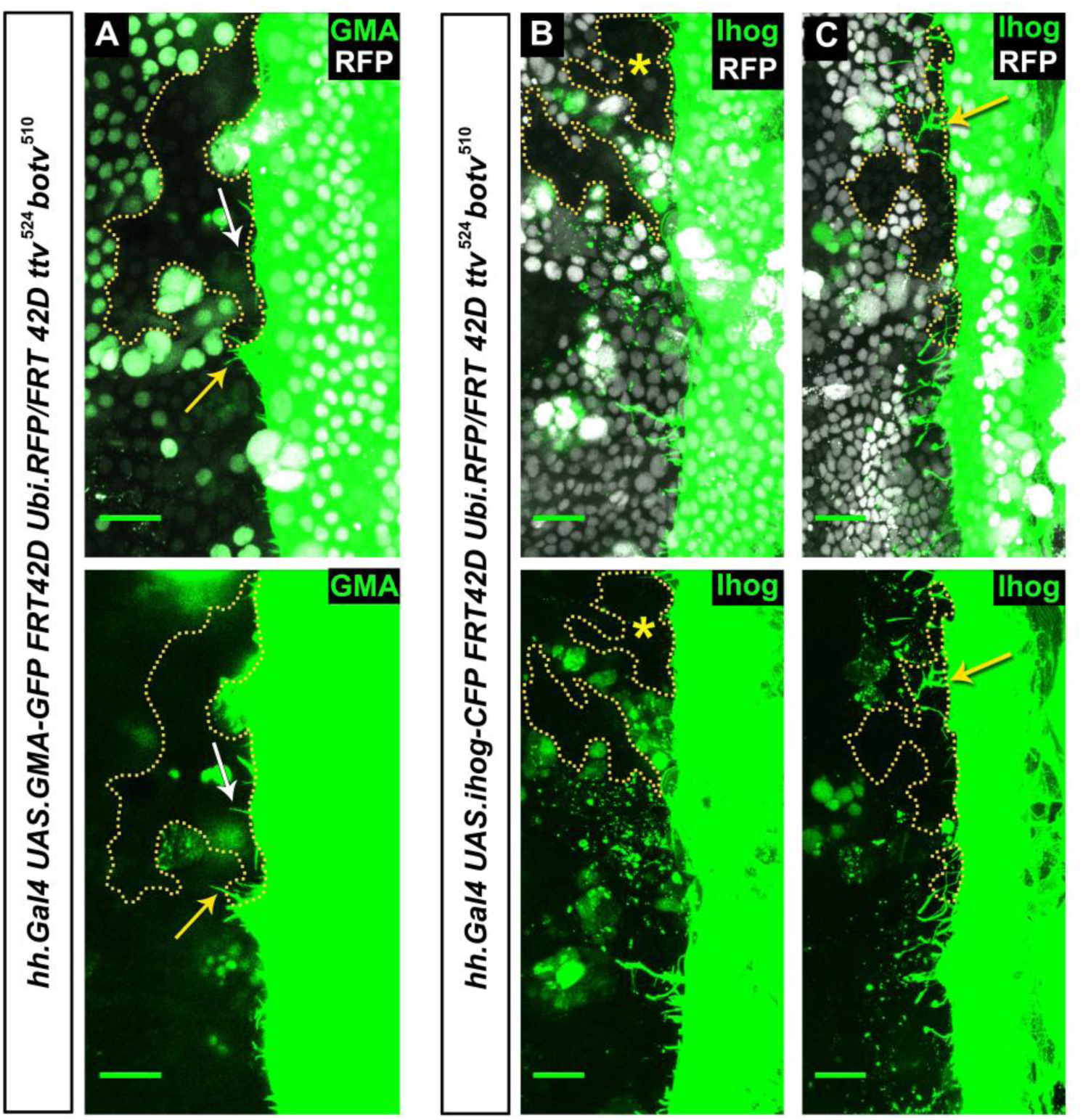
Stabilization of Ihog-labelled cytonemes by glypicans. (A-C) First frame from Movie 1 showing P compartment cytonemes crossing *ttv^524^ botv^510^* double mutant clones induced in the A compartment and abutting the P compartment (marked by the lack of RFP and delimited by a dash line). Cytonemes are labelled by GMA-GFP (A) or Ihog-CFP expression (B, C). GMA-GFP cytonemes protruding from the P compartment are able to invade the A compartment and remain dynamic (Movie 1A) independently of glypican presence (A). However, Ihog-CFP labelled cytonemes protruding from cells overexpressing Ihog (P compartment) are stabilized when they navigate wild type cells or small *ttv^524^ botv^510^* clones (C white arrow and Movie 1C), being dynamics when they navigate large *ttv^524^ botv^510^* double mutant clones areas (B asterisk and Movie 1B). Scale bar: 20 μm.

In agreement, the ectopic expression of either *UAS.ihogΔFn1-RFP* or *UAS.ihogFn1***-RFP,* which are unable to interact with Hh, noticeably affects the formation of the Hh gradient, showing a dominant negative effect on Hh sequestration in the Hh-receiving cells (Fig. 4E,F; Supplementary Fig. 5D). However, *UAS.ihogFn1***-RFP* shows a normal reception in the first cell row adjacent to the compartment border; this construct has a normal interaction with glypicans (Fig. 3G,G’), which would allow normal reception by contact between Hh presenting and receiving cells. Furthermore, the effect of both *UAS.ihogΔFn1* or *UAS.ihogFn1***-RFP* expression on the gradient is stronger than that of *UAS.ihogΔFn2-HA.* Altogether we can conclude that the Ihog Fn1 mutants, that almost abrogate the sequestration of Hh and affect the interaction with glypicans, drastically modify Hh responses and gradient formation.

### Glypicans regulate in trans the stabilization of cytonemes by ectopic Ihog expression

In physiological conditions, both Hh reception and delivery depend on dynamic cytonemes that emanate from Hh-producing and Hh-receiving cells and that present similar behaviors when analyzed using innocuous actin reporters (Bischoff *et al*., 2013; Chen *et al*., 2017; González-Méndez, Seijo-Barandiarán and Guerrero, 2017) such as the actin-binding domain of moesin fused to GFP (GMA-GFP). We had previously described that ectopic expression of Ihog stabilizes cytonemes (Bischoff *et al*., 2013; González-Méndez, Seijo-Barandiarán and Guerrero, 2017) and that such stabilization in wing disc cells requires the presence in trans (neighboring cells) of the heparan sulfate proteoglycans (HSPGs) (Bischoff *et al*., 2013) and, more specifically, of glypicans (González-Méndez, Seijo-Barandiarán and Guerrero, 2017). Thus, Ihog-glypicans interactions could be key regulators of cytoneme dynamics and their signaling function.

To test this hypothesis, we performed real time *in vivo* experiments expressing *GMA-GFP* in the P compartment of the abdominal histoblast nests and analyzed the behavior of cytonemes that elongate within *ttv^-/ -^botv^-/-^* double mutant clones in the A compartment abutting the A/P border. We observed that P compartment GMA-labelled cytonemes are able to invade the glypican-free *ttv^-/-^ botv^-/-^* mutant territory of clones touching the A/P boundary and that these cytonemes are as dynamic as those along the neighboring A compartment wild-type territory (Fig. 5A; **Movie 1A**). However, when over-expressing the wild type Ihog fused to CFP instead of GMA-GFP in the P compartment, the cytoneme stabilization effect of the ectopic expression of Ihog is absent in cytonemes navigating within large *ttv^-/-^ botv^-/-^* mutant clones in the A compartment but, as expected, present in cytonemes in the neighboring A compartment wild-type territory (Fig. 5B; **Movie 1B**) or within small *ttv^-/-^ botv^-/-^* mutant clones located close to the P compartment. We reason that, if long enough, Ihog-CFP stabilized cytonemes protruding from the P compartment cells get anchored to the wild type cell membranes expressing normal glypicans, and thus are able to reach and cross small mutant clones, but unable to cross larger glypican-deficient areas (Fig. 5C; **Movie 1C**).

Next, to assess whether this glypican requirement also has an effect on cytoneme stabilization *in cis,* we induced *ttv^-/-^ botv^-/-^* double mutant clones in the wing disc and expressed Ihog in the same cells. Ihog cytonemes emanating from glypican-deficient cells are stabilized and able to extend over wild type cells (Supplementary Fig. 6, yellow arrowhead), in contrast to cytonemes emanating from wild type cells that do not extend over glypicans deficient cells (Supplementary Fig. 4 red arrowhead). In summary, these data demonstrate that, while glypicans might not directly influence cytoneme formation, they have an effect on their dynamics *in trans*, maybe regulating Ihog presence in opposing cytonemes.

### Ihog functional extracellular domains can modulate cytoneme dynamics

To investigate the relevance of the Ihog-gypican interactions in cytoneme dynamics, we examined the Ihog domain(s) responsible for cytoneme stabilization. Cytoneme behavior was analyzed by expressing *GMA-GFP* together with either wild type Ihog or the different Ihog mutant variants in the Hh producing cells (Fig. 6A, **Movies 3, 4**) and the lifetime of cytonemes was measured. As anticipated, no cytoneme stabilization was observed after ectopic expression of the Ihog intracellular fragment (*UAS.ihogCT-RFP)* (**Movie 2C**). Complementary, we generated another transgenic line to ectopically express Ihog without the C-terminal fragment (*UAS.ihogΔCT-RFP*) and, as expected, cytoneme stabilization was observed (**Movie 2B**). Ihog cytonemes were also dynamic when co-expressing *GMA* and Ihog lacking different extracellular domains, either the Ig domains (*UAS.ihogΔIg-RFP*) (**Movie 2D**) or both FNIII domains (*UAS.ihogΔFN-RFP*) (**Movie 2E**). Likewise, the absence of either the Fn1 (**Movie 2F**) or the Fn2 domains (**Movie 2G**) results in no cytoneme stabilization. Altogether, we can conclude that Ihog ability to stabilize cytonemes resides in the extracellular domains of Ihog, including the glypican-interacting FN type III and the Ig domains.

**Figure 6.**
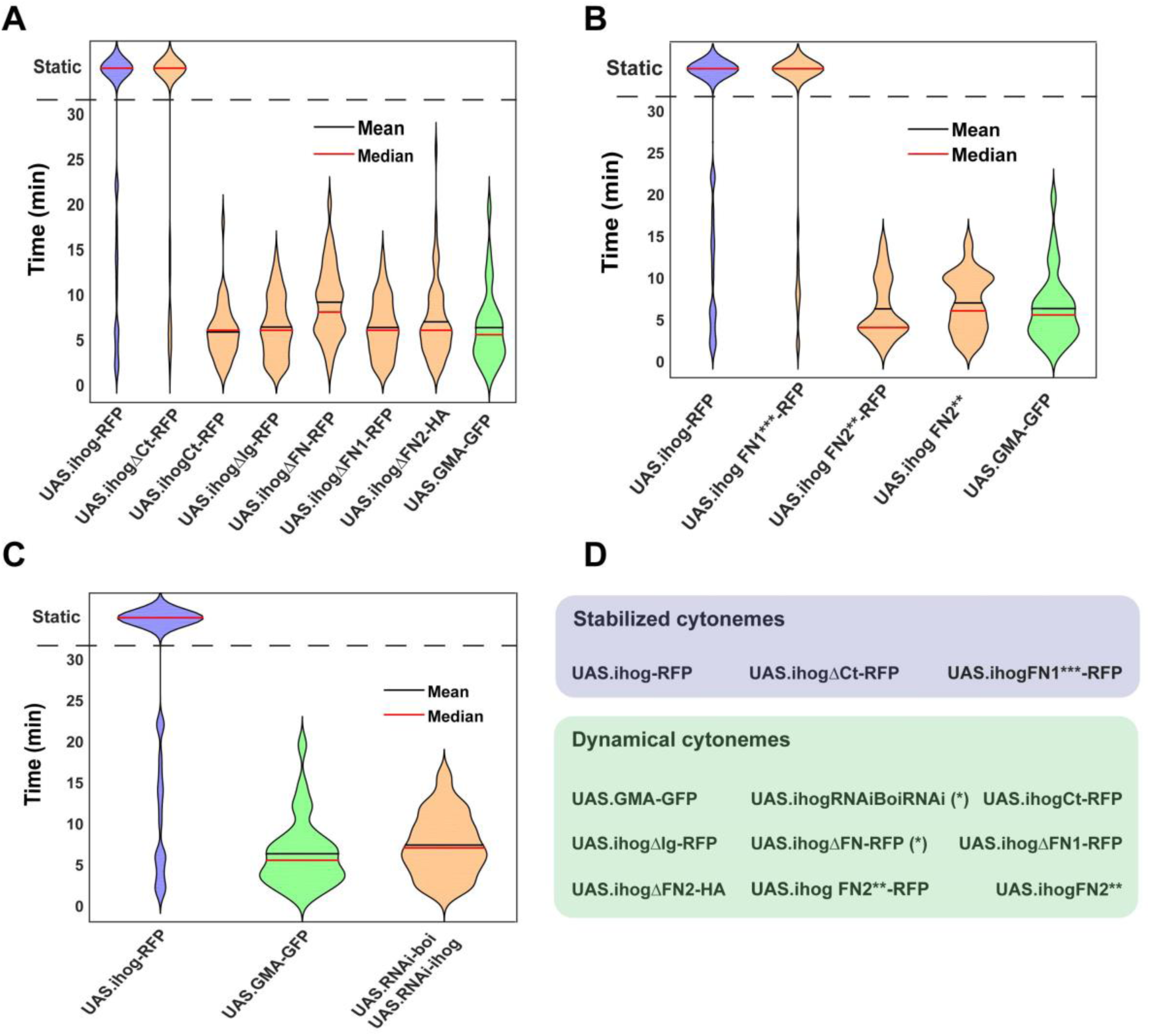
Role of Ihog domains in the regulation of cytoneme dynamics. A) Violin plots of the lifetimes distribution of dynamic cytonemes in different Ihog overexpression genotypes: *UAS.ihog-RFP* (blue), wild type control *UAS.GMA-GFP* (green), different Ihog deletion mutants (orange). B) Comparison between the effect of the ectopic expression of different Ihog point mutations: Control *UAS.ihog-RFP* (blue), wild type control *UAS.GMA-GFP* (green); mutants of Ihog (orange). C) Comparison between different levels of Ihog: *UAS.ihog-RFP* (blue), *UAS.GMA-GFP* (green), *UAS.RNAi-Boi/UAS.RNAi-ihog* (orange). D) Table summarizing the cytoneme dynamic phenotypes: Stabilized cytonemes in blue and dynamical cytonemes in green. (*) Dynamic with statistical lifetime differences from wild type *GMA-GFP* cytonemes.

Since both FNIII domains (Fn1 and Fn2) affect cytoneme dynamics, we then analyzed the effect on cytoneme stabilization of the point mutation constructs: IhogFn2**-RFP, described to modify the interaction of Ihog with Ptc (Zheng *et al*., 2010), and IhogFn1***-RFP, affecting Ihog interaction with Hh (Fig. 3C) but not with glypicans (Fig. 3G,G’). The expression of *UAS.ihogFn1***-RFP* resulted in cytoneme stabilization (Fig. 6B; **Movie 3B**), showing that the ability of Ihog to modulate cytoneme dynamics is not dependent on the same amino-acids needed for Hh binding, but rather on those involved in the interaction with glypicans. In contrast, expression of *UAS.ihogFn2\*\** with or without RFP does not stabilize cytonemes (Fig. 6B; **Movie 3C and 3D**), probably because it does not interact with Dlp and blocks the interaction with Dally (Fig. 3H,H’).

We next tested if the described effects of the ectopic expression of Ihog and the Ihog mutant forms on cytoneme dynamics in the histoblast nests can in some way be observed in the wing disc. Although both stabilized and dynamic Ihog labeled cytonemes are observed in histoblasts, in the wing imaginal disc long cytonemes are only observed in those cases in which cytonemes are stabilized in histoblasts, as happens after ectopic expression of wild type Ihog-RFP, IhogΔCT-RFP and IhogFn1***-RFP (Supplementary Fig. 7B,C,I). Therefore, in wing discs only stabilized cytonemes are visible. Consequently, in the absence of one of the Ihog FN type III domains cytonemes are not visible in the wing disc due to lack of glypican interaction. In spite of this, the expression of Ihog ΔIg in the imaginal disc results in scarce and short cytonemes (Supplementary Fig. 7E) even though its expression in the histoblast nests suggested dynamic behavior **(Movie 2D)**. This unexpected cytoneme behavior indicates that the absence of the Ig domain of Ihog might be affecting an interaction with an unknown extracellular matrix component also necessary for cytoneme stabilization.

### Excess of Ihog but not of Boi can modulate cytoneme dynamics

As Boi and Ihog have a redundant function in Hh reception (Zheng *et al*., 2010), we investigated their possible redundancy also in cytoneme stability. First, we found that ectopic expression of Boi fails to stabilize cytonemes: abdominal Hh-producing histoblasts expressing Boi emit highly dynamic cytonemes **(Movie 4)**, while co-expression of Boi and Ihog leads, as predicted, to more stable cytonemes (**Movie 5**). Coexpressing *boi* RNAi and Ihog-RFP in histoblasts we examined if Boi knock-down influences ectopic Ihog-driven cytoneme stabilization. In such condition, cells still emit stabilized cytonemes even after inhibiting Boi expression during the entire development in the P compartment (**Movie 6**). All these results indicate that although Ihog contributes to Boi build-up in stabilized cytonemes (Bilioni *et al*., 2013), this stabilization is not dependent on Boi protein levels. Therefore, Ihog induced cytoneme extension and stability is regulated by Ihog-glypican interactions and, in agreement, these extracellular matrix proteins modulate Ihog but not Boi levels (Supplementary Fig. 1).

Finally, we compared the dynamics of cytonemes after expressing GMA-GFP (**Movie 7B**), GMA-GFP and Ihog simultaneously (**Movie 7A**) or GMA-GFP in the absence of Ihog and Boi in the P compartment (**Movie 7C)**. Although ectopic Ihog labeled cytonemes are more stable, cytonemes labelled by GMA-GFP that are downregulated for Ihog and Boi have dynamics similar to those of wild type cells expressing only GMA-GFP or slightly more stable, in the limit of statistical significance (Figure. 6C). These data indicate that wild-type cells do not require these adhesion molecules to produce dynamic cytonemes.

## Discussion

### Glypicans stabilize Ihog at the plasma membranes

There is strong evidence showing that the glypicans Dally and Dlp can regulate the release, dispersal and reception of Hh ligands in *Drosophila* (for review, see (Yan and Lin, 2009; Filmus and Capurro, 2014)). On the other hand, it has previously been demonstrated that Ihog and Boi, two adhesion molecules that act as Hh coreceptors, are required for all biological responses to the Hh signaling during embryonic and imaginal development (Yao, Lum and Beachy, 2006; Camp *et al*., 2010; Zheng *et al*., 2010). Although the glypican core region of Dlp is known to function as a Hh signaling binding protein, a direct high affinity interaction between Dlp and either Hh or the Hh-Ihog complex has not been detected (Williams *et al*., 2010).

In this report we have shown sound evidence of the existence of a strong Ihog-glypican protein interaction in wing imaginal discs. First, we demonstrated that glypicans stabilize Ihog at the plasma membrane showing that its amount drastically decreases in clones double mutant for genes that encode enzymes that synthetize the HSPG (*ttv* and *btv*) and for genes encoding glypicans (*dally* and *dlp*). Under the same experimental conditions, Boi plasma membrane levels are not affected, indicating that the modulation of Ihog by glypicans is specific. The molecular mechanisms leading to this regulation are still unknown, but glypicans might be needed for either Ihog recycling process, as has been proposed for Hh (Bilioni *et al*., 2013) and Wg (Gallet, Staccini-Lavenant and Thérond, 2008) or, alternatively, for its stabilization at the plasma membrane. Second, we determined that the Ihog FNIII domains, but not the Ig domain, are the ones mediating the glypican-Ihog interaction since they are able to recruit glypicans when they are ectopically expressed. Both Ihog FNIII domains, Fn1 and Fn2, are needed for this interaction since deficiency of any of the two domains results in much lower sequestration of glypicans (Fig 7 A,B).

**Figure 7.**
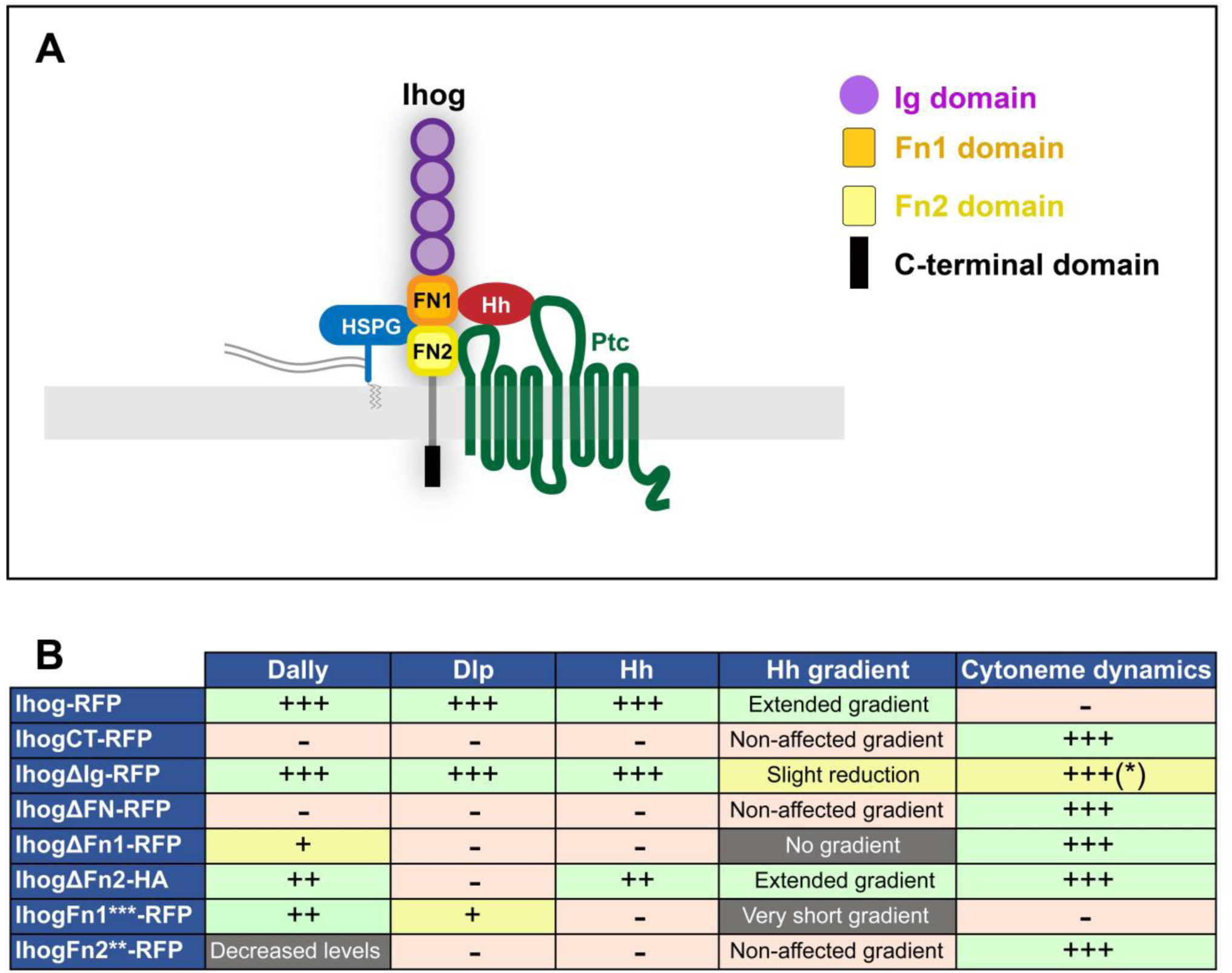
Summary of Ihog interactions with Glypicans, Hh, Ptc and table of effects induced after ectopic expression of the Ihog domains. A) Scheme depicting the Ihog domains and their inetractions of the FNIII domains with Hh (Yao, Lum and Beachy, 2006), Ptc (Zheng *et al*., 2010) and with the glypicans Dally and Dlp. B) Table summarizing the effect on Dally, Dlp, Hh gradient formation and cytoneme dynamics after ectopic expression of the different Ihog domains. Note + or - as the positive or negative effects on Dally, Dlp, Hh accretion and cytoneme dynamics. Note also by * that in the case of ectopic expression of ihogΔIg-RFP we observed that most cytonemes are dynamic in the histoblast nest, but we observed few and short stabilized cytonemes in the wing imaginal disc.

### Ihog interactions with glypicans, Hh and Ptc

Ihog has a known role in Hh recruitment to the membranes of both producing and receiving cells in the wing disc (Yan *et al*., 2010; Bilioni *et al*., 2013). It has also been proposed that Ihog may act as an exchange factor by retaining Hh on the cell surface, competing with Dlp for Hh binding (Yan *et al*., 2010). Finally, of the three Ihog extracellular domains (Fn1, Fn2 and Ig), only Fn1 and Fn2 are necessary for *in vivo* Hh signaling, having independent functions (McLellan *et al*., 2006; Zheng *et al*., 2010).

The Fn1 domain has been described as specifically needed for Hh binding (McLellan *et al*., 2006; Yao, Lum and Beachy, 2006) (Fig. 7A). In agreement, we have observed that the ectopic expression of a variant of Ihog lacking Fn1 does not accumulate Hh in the P compartment as the wild type Ihog does. The Ihog amino-acids mediating the interaction with Hh have been also identified on the Fn1 extracellular domain in tissue culture cells (Wu *et al*., 2019). The amino-acids described (Wu *et al*., 2019) are not the same as the ones identified here, although they are in the same protein region. Alteration of the amino-acids D558N, N559S, E561Q in the Fn1 domain (Ihog Fn1***) impedes Hh build-up. Interestingly, these amino-acid alterations maintain the Ihog-glypican interaction, showing that these two interactions (with Hh and glypicans) are mediated by different sequences in the same Ihog extracellular domain. Accordingly, crystal structure of a Hh-binding fragment of Ihog showed that heparin binding induces the Ihog dimerization required to mediate high-affinity interactions between Ihog and Hh; again, the Fn1 domain is specifically needed for this heparin-dependent binding. The putative heparin-binding site near the Ihog-Ihog dimer interface was found to be separated from the basic strip bridging Hh to IhogFn1 (McLellan *et al*., 2006). However, the expression of a construct only carrying the wild type Fn1 reveals that Fn1 alone recruits very low Hh levels. This low Hh recruitment suggests that although Fn1 interacts physically with Hh, the cooperation between Fn1 and Fn2 domains is needed for an efficient Fn1-Hh binding. In agreement, both domains are required for proper Ihog-glypican interaction. This result also explains that Fn2 domain influences the Hh accumulation.

In addition, the expression of either Ihog ΔFn1 or Ihog Fn1*** in the receiving cells blocks the sequestration of Hh during reception, showing that these Ihog mutant forms behave as dominant negative for this trait. This effect is observed when overexpressing most of the Ihog mutant forms, but it is particularly strong in the case of IhogΔFn1. It is possible that the association of endogenous Ihog and Boi needed for Hh reception may also be affected (Zheng *et al*., 2010).

As coreceptors, Ihog and Boi have a crucial role in the Hh high-affinity binding to Ptc. An Ihog Fn2-mediated surface presentation of Ptc has been proposed to increase either its transport rate to the cell surface or the duration of its presence (Zheng *et al*., 2010); so far, there are not enough evidences to distinguish between these two possibilities. In favor of Fn2 Ihog-mediated surface presentation of Ptc, we have shown here that the ectopic expression of Ihog-ΔFn2-HA in the receiving cells flattens the Hh gradient. Similar flattening effect has been described after abrogation of the Ptc vesicular trafficking required for its presentation at the plasma membrane to interact with Hh (González-Méndez *et al*., 2020). In our hands, however, ectopic expression of the Ihog-Fn2** construct mutated in the amino-acids thought to be required for its interaction with Ptc (K653 and Q655) (Zheng *et al*., 2010) did not accumulate Hh and Dlp and did not show an effect on the Hh gradient. All these behaviors were surprising since the deletion of the entire Fn2 domain results in Hh accretion, flattening of the Hh gradient and slight build-up of glypicans, indicating that probably the Ihog-Fn2** protein has an anomalous function (a neomorph in the classical terminology). It is possible that these two-point mutations in Fn2 somehow modify Ihog protein structure, not only avoiding its interaction with Ptc but also acting as a dominant negative in the interaction with Dally, although without affecting the Hh gradient.

### Ihog localization at cytonemes requires glypicans

Hh graded distribution across the receiving *Drosophila* epithelia is mediated by cytonemes (Bischoff *et al*., 2013) and Hh reception seems to happen at discrete contact sites along presenting and receiving cytonemes. These contact places, where most of the extracellular components needed for Hh signaling colocalize (González-Méndez, Seijo-Barandiarán and Guerrero, 2017; González-Méndez *et al*., 2020), suggest a contact-dependent mechanism for Hh reception in which receptor, coreceptors and ligand are present in close proximity in a dynamic manner, a mechanism previously described as a synaptic-like process (Huang, Liu and Kornberg, 2019).

Since cytonemes are dynamic structures that extend and retract with a defined velocity (González-Méndez, Seijo-Barandiarán and Guerrero, 2017), we analyzed the influence of glypicans, Ihog and Boi in cytoneme behavior. We had previously described that in wing discs and in the abdominal histoblast nests the ectopic expression of Ihog stabilizes cytonemes, although without affecting their normal length (Bischoff *et al*., 2013; González-Méndez, Seijo-Barandiarán and Guerrero, 2017). In the wing disc, this stabilization requires the presence in trans (neighboring cells) of the HSPG (Bischoff *et al*., 2013) and, more specifically, of glypicans (González-Méndez, Seijo-Barandiarán and Guerrero, 2017). We have demonstrated here that: 1) only Ihog but not Boi has the cytoneme stabilizing function *in vivo*; 2) the levels of glypicans are key to maintain Ihog, but not Boi, protein levels at cytoneme membranes; 3) glypicans are needed *in trans* for cytoneme stabilization; and 4) the interactions that regulate this stabilization are mediated by the Ihog FNIII domains, which are also involved in the recruitment at cytoneme contacts of Hh ligand, the reception components and the glypicans.

Even though we have observed that the simultaneous absence of both glypicans has little effect in the formation and dynamics of GMA-GFP labelled cytonemes, Ihog-stabilized cytonemes cannot cross *ttv^-/-^ botv^-/-^* mutant clones, because these cytonemes cannot extend when encountering HSPG deficient areas (as previously observed in the wing disc (Bischoff *et al*., 2013)). In contrast, Ihog-labeled cytonemes protruding from *ttv^-/-^ botv^-/-^* double mutant cells can extend and navigate over cells with normal HSPG content. We can then conclude that the interaction *in trans* of the ectopic Ihog with glypicans is essential for the effect of Ihog on stabilization of cytonemes. This mutual need is also shown through the expression of Ihog mutants for either Fn1 or Fn2 domains that do not interact well with glypicans (Fig. 7A,B)

It has been proposed that Ihog and glypican expression patterns in the wing disc could indicate a functional stoichiometric relationship. Thus, Ihog and Dally are downregulated and Dlp is upregulated at the A/P compartment border (Zheng *et al*., no date; Yan *et al*., 2010; Bilioni *et al*., 2013; Hsia *et al*., 2017), what might be necessary for proper cytoneme behavior and Hh release or reception. In agreement, it was previously described that, in the Hh producing cells, *dlp* loss of function increases the retention of Hh at plasma membranes (Callejo *et al*., 2011) while loss of function of either *dally* or *ihog* decreases this retention (Yao, Lum and Beachy, 2006; Yan *et al*., 2010; Bilioni *et al*., 2013). Since Ihog strongly attaches Hh to cytonemes, glypicans could aid Hh maintenance at cytonemes, its presentation to Ptc and its release during the reception process. Accordingly, we have demonstrated that ectopic Ihog cannot stabilize cytonemes when the surrounding cells are mutant for glypicans or when Ihog is mutated in the domains that promote its interaction with glypicans. Intriguingly, the same Ihog domains that regulate cytoneme dynamics are those also involved in the recruitment of Hh ligand, glypicans and the reception complex. It was previously demonstrated that this complex is localized at discrete contact sides along cytonemes (González-Méndez, Seijo-Barandiarán and Guerrero, 2017). Our results have also reinforced the previously described autonomous requirement of Dlp in Hh reception (Desbordes and Sanson, 2003; Lum, Yao, *et al*., 2003; C. Han *et al*., 2004; Takei *et al*., 2004; Williams *et al*., 2010). Based on their interaction with Ihog, we propose that glypicans play a role in the localization of the receptor complex in cytonemes.

Glypicans regulate several morphogenetic gradients in addition to that of Hh (ie.Wnt, Bmp, FGF). It has also been described that Dlp class of glypicans shields the highly hydrophobic lipid moiety of Wnts from the aqueous environment helping to be handed over to Wnt receptors (McGough *et al*., 2020). Dlp could hide the lipids while transferring Hh to its receptor Ptc. However, how glypicans specifically control these pathways was unknown. We show here that glypicans regulate Hh signaling through the interaction with its coreceptor Ihog regulating its presence at the plasma membrane. This interaction could then confer the specificity of glypicans for Hh signaling.

## Material and Methods

### Generation of the Ihog constructs

IhogΔCT was obtained by PCR amplification using the Ihog cDNA as a template (*5’CACCATGGACGCTGCTCACATCCTC3’;5’CTTGTTTGGATTGTTTCCCCGGCTTC3’*). The resulting product was then cloned into the entry vector pENTR/D-TOPO by directional TOPO cloning (Gateway system; Invitrogen), and introduced by recombination into the destination vector pTWR (pUAST-RFP).

IhogΔFN construct was obtained by PCR amplification using the Ihog/pENTR vector as a template (*5’GAATTTAGTGCCCTTAAGCAGG3’; 5’CTGCTTCTGCCTGGAACCCT3’*). The resulting product was then ligated obtaining the IhogDFN/pENTR vector, lacking the FN domains, that was then introduced by recombination into the destination vector pTWR (pUAST-RFP).

To generate the IhogΔIG, IhogΔFn1, IhogΔFn2 and the IhogCT constructs two different PCR reactions were performed (primers), introducing a restriction site (BamHI, NdeI or KpnI). After restriction and subsequent ligation, the product lacking the domain of interest was cloned into the entry vector pENTR/D-TOPO by directional TOPO cloning (Gateway system; Invitrogen) and introduced by recombination into the destination vector pTWR (pUAST-RFP) or pTWHA (pUAST-RFP).

Primers:

*ΔIG*

*PCR1:*

*5’CACCATGGACGCTGCTCACATCCTC3’;5’CACCAAAGATTC<u>GGATCC</u>GCGAAG3’ PCR2: 5’CAGATTCAGGGATCCCGGGAATCG3’;5’CACGCCAACGCTGTTGAGGCT3’*

*ΔFN1*

*PCR1:*

*5’CACCATGGACGCTGCTCACATCCTC3’;5’CTTCTGCCT<u>GGTACC</u>CTGATTGGGC3’ PCR2:5’GCAGCCAGGTACCGCACTTGATCCG3’;5’CACGCCAACGCTGTTGAGGCT3’*

*ΔFN2*

*PCR1:5’CACCATGGACGCTGCTCACATCCTC3’;5’GTATTCCTCGATCTC<u>CATATG</u>CTCT GGCAC3’ PCR2:5’GAATTTAGTGCCCATATGCAGGGG3’;5’CACGCCAACGCTGTTGAGGCT3’*

*CT*

*PCR1:5’CACCATGGACGCTGCTCACATCCTC3’; 5’CACCAAAGATTC<u>GGATCC</u>GCGAAG3’ PCR2:5’CGCACCCAAGGATCCAAAACCAGC3’;5’CACGCCAACGCTGTTGAGGCT3’*

To generate the construct IhogFN1***, carrying point mutations D558N, N559S and E561Q, a PCR was performed with the following primers: 5’TGAAGTTG<u>GAG</u>TGTAAGGCCA3’ and 5’TCAAGTTC<u>AAC</u>GT<u>CAG</u>GAAGTCA3’. pTWR-IhogRFP vector was digested using BglII and XhoI restriction enzymes. The PCR product and the digested vector were put together by Gibson Assembly.

The construct IhogFN2** (point mutations K653E and Q655E) was generated using pTWR-IhogRFP as a template. First, a mutagenized fragment was generated by exchanging the first nucleotide for guanosine in the codons of K653 and Q655 (Zheng *et al*., 2010). The mutagenized fragment was then inserted into pTWR-IhogRFP by digestion with Xho1 and NgoMIV.

The construct Fn1-CD8mCherry was generated by amplification of FN1 domain with the primers: 5’AGGCATGCATCAAGTGCTGCACCTG3’ and 5’ACGGTACCCCACCGACTCCTCCAAATG3’. The fragment was then cloned by restriction ligation into the Kpn1 Sph1 sites of the pLOT-CD8-mCherry vector (Harmansa *et al*., 2015).

### Drosophila strains

The description of mutations, insertions and transgenes is available at Fly Base (http://flybase.org). The following mutants and transgenic strains were used: *tub.Gal80^ts^*, *hs-Flp122* (Bloomington Drosophila Stock Centre (BDSC), Indiana, USA; http://flystocks.bio.indiana.edu), *hs-FLP* (Golic and Lindquist, 1989), *dally^32^* (Franch-Marro *et al*., 2005), *dlp^20^* (Franch-Marro *et al*., 2005), *ttv^524^* (Takei *et al*., 2004), *botv^510^* (Takei *et al*., 2004), *boi* (Zheng *et al*., 2010) and *ihog^Z23^* (Zheng *et al*., 2010). The reporter lines used were: *Bac Hh:GFP* (Chen *et al*., 2017), *Bac ihog:GFP* (Hsia *et al*., 2017), and *dally-trap-YFP* (Lowe *et al*., 2014).

### Overexpression experiment

The following Gal4 and LexA drivers were used for ectopic expression experiments using Gal4/UAS (Brand and Perrimon, 1993) and LexA/LexAop (Yagi, Mayer and Basler, 2010) systems: *hh.Gal4* (Tanimoto *et al*., 2000), *ptc.Gal4* (Hinz, Giebel and Campos-Ortega, 1994), *ap.Gal4* (Calleja *et al*., 1996), *ap.LexA* and *hh.LexA*. The transgene act>y+>Gal4 (Pignoni and Zipursky, 1997) was used to generate random ectopic clones of the UAS lines. Larvae of the corresponding genotypes were incubated at 37°C for 12 minutes to induce hs-Flp-mediated recombinant clones. We use the tub-Gal80^ts^ for the transient expression of transgenic constructs with the UAS/Gal4 system. Fly crosses were maintained at 17°C and dissected after inactivation of the Gal80^ts^ repressor for 24 hours at restrictive temperature (29°C).

The UAS-transgenes used were: *UAS.ihog-RFP* (Callejo *et al*., 2011), *UAS.ihog-CFP* (Bilioni *et al*., 2013), *UAS.ihogΔCT-RFP*, *UAS.ihogΔFN-RFP*, *UAS.ihogΔIg-RFP UAS.ihogΔFn1-RFP*, *UAS.ihogΔFn2-HA*, *UAS.ihogFn1***-RFP*, *UAS.ihogFn2**-RFP, UAS.ihogCT-RFP*, *UAS.boi-YFP* (Bilioni *et al*., 2013), *UAS.GMA-GFP* (Bloor and Kiehart, 2001), *UAS.ihog-RNAi* (VDRC 102602), *UAS.boi-RNAi* (VDRC 108265), *UAS.dlp-RNAi* (VDRC 10299), *UAS.dally-RNAi* (VDRC 14136). *UAS.ihogFn2\*\** (Zheng *et al*., 2010), referred as *UAS.ihogFN2\** in their study) were also used as controls to test whether or not they gave the same results as the expression of the same Ihog RFP-tagged forms and certainly they gave the same phenotypes in all the experiments.

### Clonal analysis

Mutant clones were generated by heat shock Flp-mediated mitotic recombination. For wing imaginal disc dissection, individuals grown at 17°C were incubated at 37°C for 45 minutes 48–72 hours after egg laying (AEL). The genotypes were:

− *y w* hsFlp^122^/BacihogGFP; *dally^32^ dlp^20^* FRT2A / Ubi.RFP FRT2A
− *y w* hsFlp^122^/BacihogGFP; *dally^32^* FRT2A / Ubi.GFP FRT2A
− *y w* hsFlp^122^/BacihogGFP; *dlp^20^* FRT2A / Ubi.GFP FRT2A
− *y w* hsFlp^122^, *boi*/Y; *ihog^Z23^* FRT40A / Ubi.RFP FRT40
− *y w* hsFlp^122^, *boi*/Y; *ihog^Z23^* FRT 40A / Ubi.RFP FRT40; *dally trap*-YFP

For *in vivo* movies, individuals were incubated twice at 37°C for 45 minutes with a 24-hours interval during L3. The genotypes were:

− *y w* hsFlp^122^; FRT42D *ttv^524^ botv^510^*/FRT42DUbi.RFP; *UAS.GMA-GFP*/*hh.Gal4, tub*Gal80^ts^.
− *y w* hsFlp^122^; FRT42D *ttv^524^ botv^510^*/FRT42D Ubi.RFP; *UAS.ihog-CFP*/*hh.Gal4*, *tub*Gal80^ts^.

### Immunostaining of imaginal discs

Immunostaining was performed according to standard protocols (Capdevila and Guerrero, 1994). Imaginal discs from third instar larvae were fixed in 4% paraformaldehyde in PBS for 20 minutes at room temperature (RT) and permeabilized in PBS 0,1% Triton (PBT) before incubating with PBT 1% BSA for blocking (1 hour at RT) and then with the primary antibody (overnight at 4°C). Three washes at RT for 15 minutes and incubation with the appropriate fluorescent secondary antibodies (ThermoFisher) at a 1:400 dilution for 1 hour at room temperature and then washing and mounting in mounting media (Vectashield). Primary antibodies were used at the following dilutions: mouse monoclonal α-Ptc ((Capdevila and Guerrero, 1994); Hybridoma Bank Iowa), 1:30; mouse monoclonal α-Dlp ((Lum, Yao, *et al*., 2003); Hybridoma Bank, Iowa), 1:30; rabbit polyclonal α-βGal (from Jackson laboratories), 1:1000; rabbit polyclonal α-GFP (Molecular Probes, α-6455), 1/1000, rabbit anti HA polyclonal antibody (SIGMA), 1:100; rabbit polyclonal α-Ihog (Bilioni *et al*., 2013), 1:100; mouse monoclonal α-En (Patel *et al*., 1989), 1:100; and rat monoclonal antibody α-Ci ((Motzny and Holmgren, 1995) a gift from B. Holmgren), 1:20.

### Microscopy and image processing of wing imaginal discs

Laser scanning confocal microscope (LSM710 Zeiss) was used for confocal fluorescence imaging of wing imaginal discs. ImageJ software (National Institutes of Health) was used for image processing and for image analysis. The number (n) of analyzed wing discs in each experiment and for each genotype was: Fig. 2, n>5; Fig. 3 and Supplementary fig. 3 (Hh retention), n>4; Fig. 3 (glypicans interaction), n>5; Fig. 4, n>7; Supplementary fig. 1 (Ihog reduction and Boi levels), n>10 for each genotype; Supplementary fig.4, n>4; Supplementary fig. 5, n>4; Supplementary Fig. 7, n>5.

Profiles of Ihog, Boi, Dlp, Dally and Hh were made taken intensity gray values of a dorso-ventral region of the wing disc and normalizing to the mean value gray value of the ventral compartment.

Profiles of Ptc were obtained taken intensity gray values of an anterior-posterior region in both ventral (experimental data) and dorsal compartment (control data). For the normalization of each profile we subtracted the background using the average value of the posterior compartment, and then normalizing the data to the minimum value of the anterior compartment.

### *In vivo* imaging of pupal abdominal histoblast nests

Imaging of pupal abdominal histoblasts was done using a chamber as described in (Seijo-Barandiarán, Guerrero and Bischoff, 2015). Hh signaling filopodia from histoblasts of dorsal abdominal segment A2 were filmed using 40x magnification taking Z-stacks of around 30 mm of thickness with a step size of 1 mm every 1 or 2 minutes, depending on the experiment, using a LSM710 confocal microscope. All movies were analyzed with Fiji. All imaged pupae developed into pharate adults and hatched normally.

### Quantification method and numerical analysis of cytoneme dynamics

To statistically compare the cytoneme lifetime when *boi* and *ihog* expression is altered, we used the method described in (González-Méndez, Seijo-Barandiarán and Guerrero, 2017), tracking cytonemes with the GMA-GFP signal (*hh.Gal4 tubG80ts UAS.ihog-RFP UAS.GMA-GFP* n=4; *hh.Gal4 UAS.GMA-GFP* n=6; *hh.Gal4 UAS.boi-RNAi UAS.ihog.RNAi UAS.GMA-GFP* n=7).

To statistically compare the cytoneme lifetime when the Ihog proteins, either the wild type form or the mutant forms, are expressed ectopically, we manually quantified the frames in which cytonemes have been observed tracking the GMA-GFP signal. For each condition we scored 3 to 5 pupae and between 10 and 20 cytonemes per pupa (*hh.Gal4 tubG80ts UAS.ihogΔCt-RFP/UAS.GMA-GFP* n=3; *hh.Gal4 tubG80ts UAS.ihogCt-RFP/UAS.GMA-GFP* n=4; *hh.Gal4 tubG80ts UAS.ihogΔIg-RFP*/*UAS.GMA-GFP* n=5; *hh.Gal4 tubG80ts UAS.ihogΔFN-RFP/UAS.GMA-GFP* n=5; *hh.Gal4 tubG80ts UAS.ihogΔFN1-RFP*/ *UAS.GMA-GFP* n=6; *hh.Gal4 tubG80ts UAS.ihogΔFN2-HA/UAS.GMA-GFP* n=4; *hh.Gal4 tubG80ts UAS.ihogFN1***-RFP*/*UAS.GMA-GFP* n=4; *hh.Gal4 tubG80ts UAS.ihogFN2**-RFP/UAS.GMA-GFP* n=5; *hh.Gal4 tubG80ts UAS.ihogFN2**/UAS.GMA-GFP* n=5).

The manual quantification of the data was stored in an excel file and uploaded in a Matlab script. The Matlab script was designed to organize the data, to compute the statistical analysis and to represent the results in different violin plots (Figure 6). For genotypes showing no static (only dynamic) cytonemes, the statistical analysis was done using their numerical lifetimes. To examine the normality of the data distribution, we performed a Shapiro-Wilk test. Since the results showed a non-parametric distribution of the experimental data, we selected a Wilcoxon rank sum test to compare the numerical lifetimes between two genotypes. The resulting p-values can be observed in Supplementary Tables 1 and 2).

For genotypes showing static and no static cytonemes we defined a no numerical case to quantify the frequency of static cytonemes. As a result, we obtained a “mixture” violin plots representing the distribution of the lifetime of the whole cytoneme population.

**Supplementary.T.1.** Statistical analysis of the lifetime for different Ihog levels. P-values obtained from Wilcoxon rank sum test to statistically compare cytoneme lifetimes for different levels of Ihog (n.s not significance in grey, significance with the correspond p value in scientific notation in orange).

| Wilcoxon rank sum test | UAS.ihog-RFP | UAS.GMA-GFP | UAS.RNAi-boi;UAS.RNAi-ihog |
| --- | --- | --- | --- |
| UAS.ihog-RFP |  | n.s | n.s |
| UAS.GMA-GFP |  |  | 5,00E-02 |
| UAS.RNAi-boi; UAS.RNAi-ihog |  |  |  |

**Supplementary.T.2.**
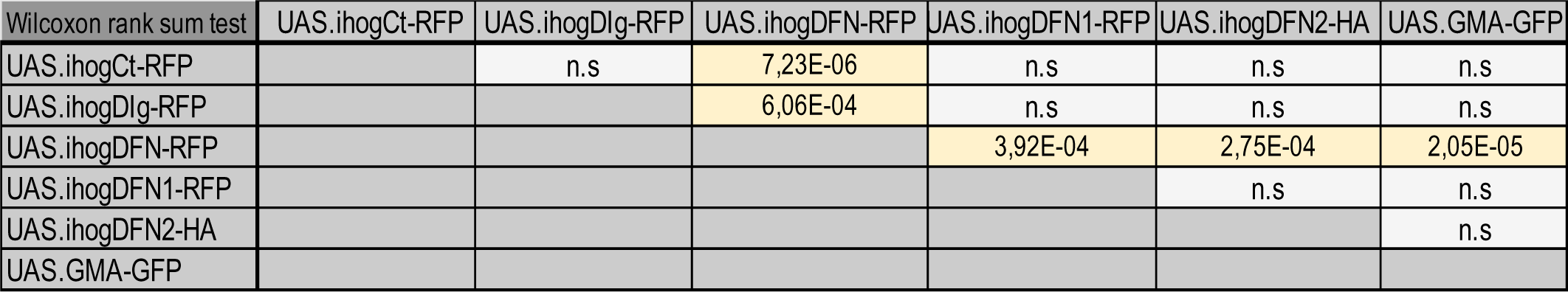
Statistical analysis of the lifetime for different Ihog mutants. P-values obtained from Wilcoxon rank sum test to statistically compare cytoneme lifetimes for different Ihog mutants with the control (UAS.GMA-GFP) (grey: n.s= not significance, orange: significance with the correspond p-value in scientific notation).

| Wilcoxon rank sum test | UAS.ihogCt-RFP | UAS.ihogDlg-RFP | UAS.ihogDFN-RFP | UAS.ihogDFN1-RFP | UAS.ihogDFN2-HA | UAS.GMA-GFP |
| --- | --- | --- | --- | --- | --- | --- |
| UAS.ihogCt-RFP |  | n.s | 7,23E-06 | n.s | n.s | n.s |
| UAS.ihogDlg-RFP |  |  | 6,06E-04 | n.s | n.s | n.s |
| UAS.ihogDFN-RFP |  |  |  | 3,92E-04 | 2,75E-04 | 2,05E-05 |
| UAS.ihogDFN1-RFP |  |  |  |  | n.s | n.s |
| UAS.ihogDFN2-HA |  |  |  |  |  | n.s |
| UAS.GMA-GFP |  |  |  |  |  |  |

### Western-blot analysis

The expression levels of the proteins induced by the UAS constructs were analyzed by western blotting (Supplementary Fig 2). Protein extracts from third instar larvae of *tubGal4/tubGal80^ts^;UAS.ihog.RFP/+*, *tubGal4/tubGal80^ts^;UAS.ihogCT-RFP/+, tubGal4/tubGal80^ts^;UAS.ihogΔIg-RFP/+,tubGal4/tubGal80^ts^;UAS.ihogΔFN-RFP/+*, *tubGal4/tubGal80^ts^;UAS.ihogΔFn1-RFP/+* and *tubGal4/tubGal80^ts^;UAS.ihogΔFn2-HA/+* genotypes were prepared in lysis buffer containing protease inhibitors. Samples were resolved by SDS-PAGE, immunoblotted, and incubated with rabbit anti-Ihog 1:500 antibody (Bilioni et al., 2013) (SDIX). The samples were resuspended in sample buffer with β-mercaptoethanol and subjected to 1 D SDS-Page (8%) and Western blotting Blotted membranes were probed with appropriate antibodies, either to HA (1:500 mouse monoclonal, Covance) or RFP (1:5000 rabbit polyclonal Chromoteck). Blots were incubated with fluorescent α-mouse (800CW) and α-rabbit (680RD) secondary antibodies (Li-Cor) and imaged with the 364-Odyssey equipment.

## Supporting information

Movie 1. Cytoneme stabilization by ectopic Ihog require Glypican function in the neighboring cells.

Movie 2. Roles of Ihog domains on cytoneme stability

Movie 3. Different effect of Ihog-Hh and Ihog-Ptc interaction on cytoneme stability

Movie 4. Cytoneme dynamics of abdominal histoblasts expressing UAS.boi-YFP

Movie 5. Cytoneme dynamics of histoblasts expressing both UAS.boi-YFP and UAS.ihog-RFP

Movie 6. Cytoneme dynamics of histoblast expressing UAS.ihog-RFP and UAS-RNAi-boi

Movie 7. Different levels of Ihog and Boi in histoblasts affect cytoneme dynamics

## Acknowledgements

We are grateful to Ana-Citlali Gradilla for invaluable advice and comments on the manuscript and to Marcus Bischoff for his expertise and advice in setting up the in vivo imaging of the abdominal histoblasts in our lab and for hosting I.S-B to learn to do it. We thank T. Kornberg, X. Zheng, P. Beachy, M. Bischoff, JP Vincent and R. Holmgren for stocks and reagents and to X.Zheng for her help in the characterization of Ihog mutant constructs. We also thanks to Transgenesis and Confocal Facilities of the CBMSO and to Bloomington and Vienna stock centers for fly stocks. This work was supported by grants BFU2014-59438-P and BFU2017-83789-P and TENTACLES consortium RED2018-102411-T to IG from the Spanish Ministry of Science and Innovation and University and by institutional grants from the Fundación Areces and from Banco de Santander to the CBMSO. FPI fellowship from the Spanish Ministry of Science and Innovation and University supported C.J. (BFU2017-83789-P), A.A-T (BFU2014-59438-P), and I.S-B. (BFU2011-25987). ‘Fellowships for Excellence’ from the International PhD Program in Molecular Life Sciences of the Biozentrum, University of Basel supported GA.

---

**Supplementary Figure 1.**
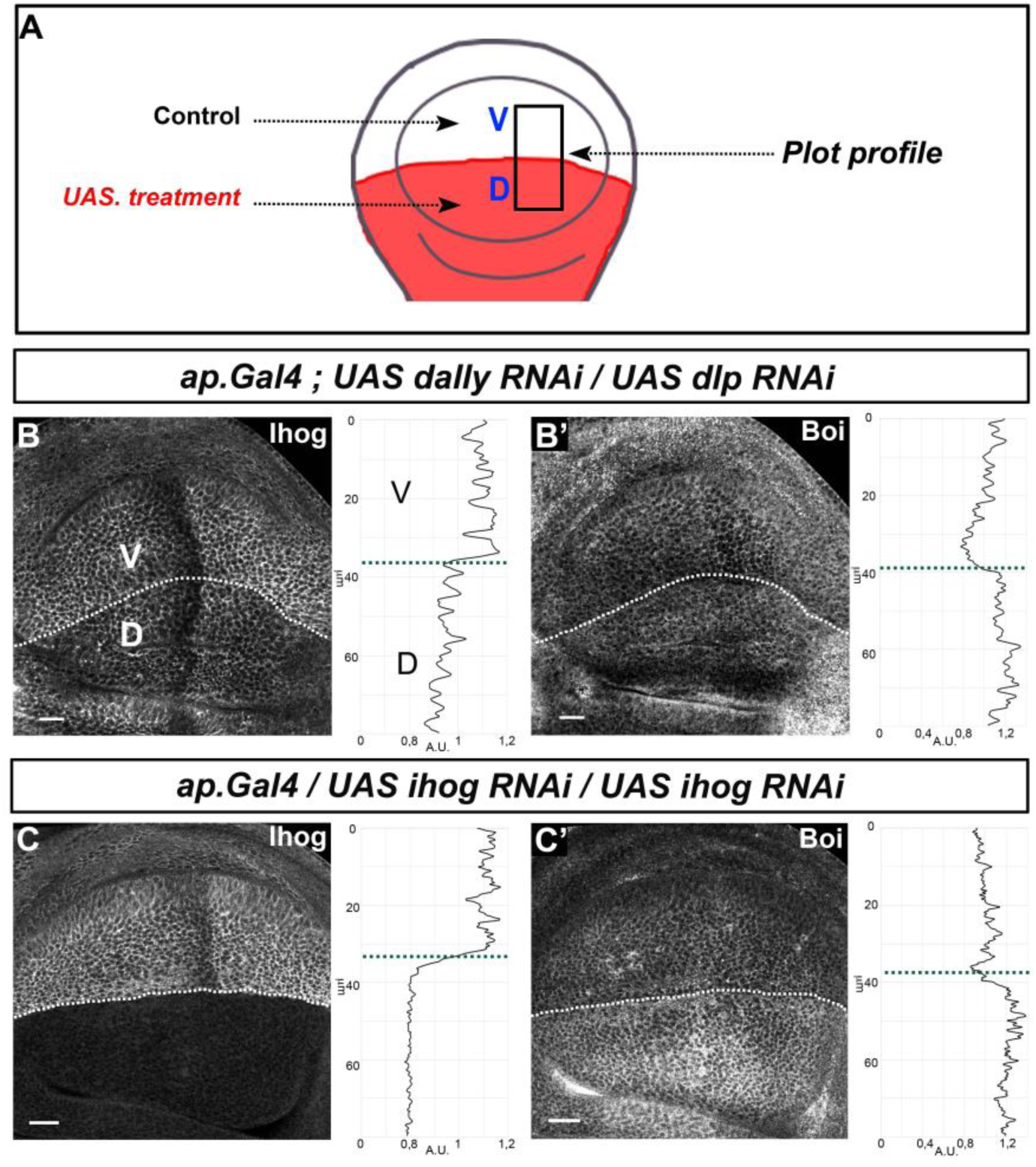
Glypicans modulate the levels of Ihog but not of Boi. A) Scheme to depict the *ap.Gal4* expression domain in the ventral part of the wing disc. B) Ihog expression of *BacIhog-GFP* in *apGal4/+; UAS dally-RNAi/+* and *apGal4/+; UAS dlp-RNAi/+* wing discs. B’) Boi expression using anti-Boi antibody in *apGal4/+; UAS dally-RNAi/+* and *apGal4/+; UAS dlp-RNAi/+* wing discs. C) Ihog expression of *BacIhog-GFP* in *apGal4/+; UAS ihog-RNAi/+* wing discs. C’) Boi expression using anti-Boi antibody in *apGal4/+; UAS ihog-RNAi/+* wing discs. Plot profiles were performed over a ROI of size 80μm X 30μm. Note the decrease of Ihog expression in the dorsal compartment compared to the wild type ventral compartment after knocking down Dally and Dlp. Note also that Boi expression is slightly increased in the same mutant conditions and strongly upregulated after Ihog elimination by RNAi expression. Scale bar: 20 μm. Discs are oriented with the dorsal part down, ventral up and posterior right.

**Supplementary Figure 2.**
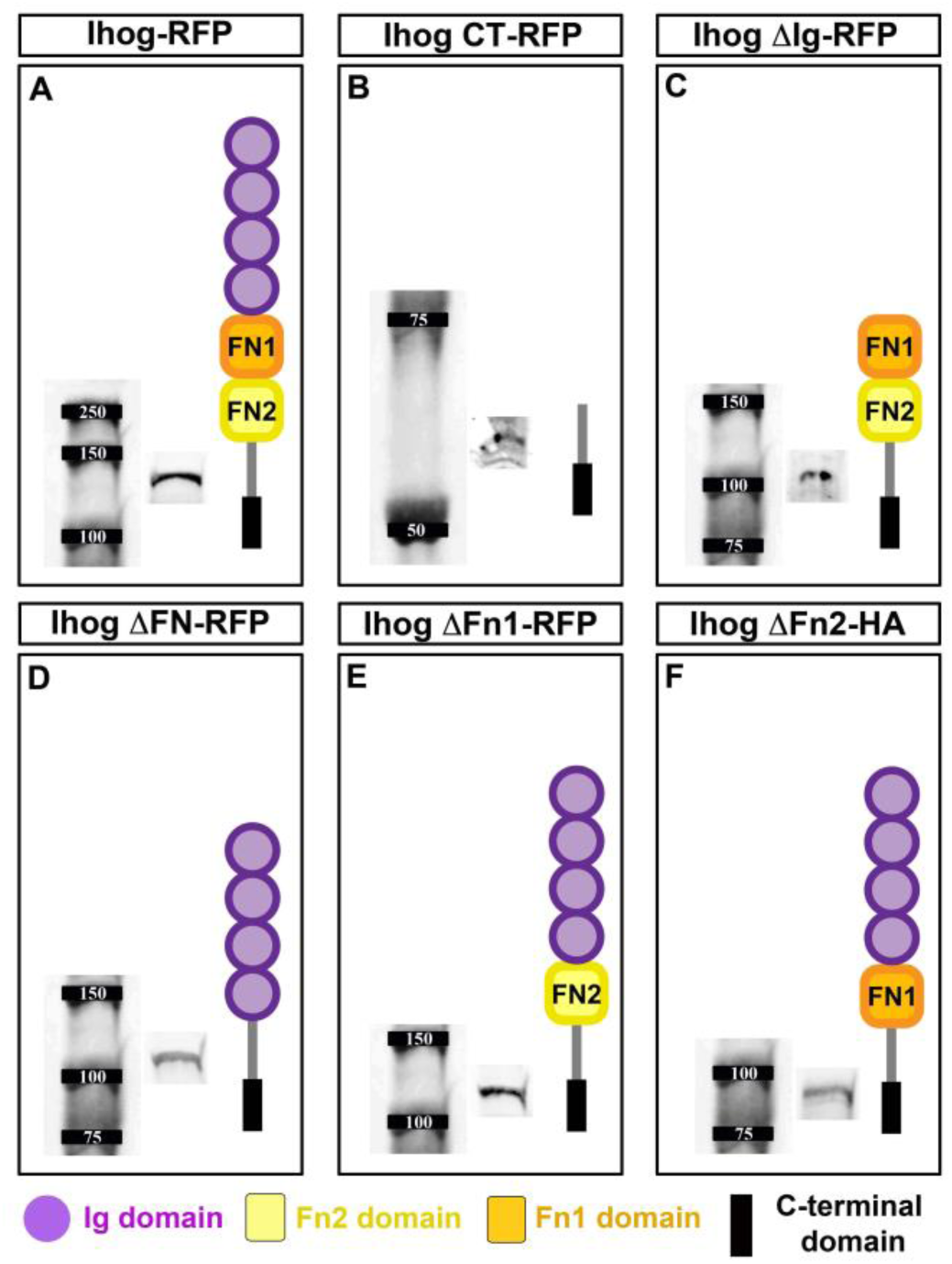
Ihog mutant constructs. (A-F) scheme of the different Ihog mutant forms: Ihog-RFP (A); IhogCT-RFP (B); IhogΔIg-RFP (C); IhogΔFN-RFP (D); IhogΔFn1-RFP (E); IhogΔFn2-HA (F) and their molecular weight shown by Western blot analysis.

**Supplementary Figure 3.**
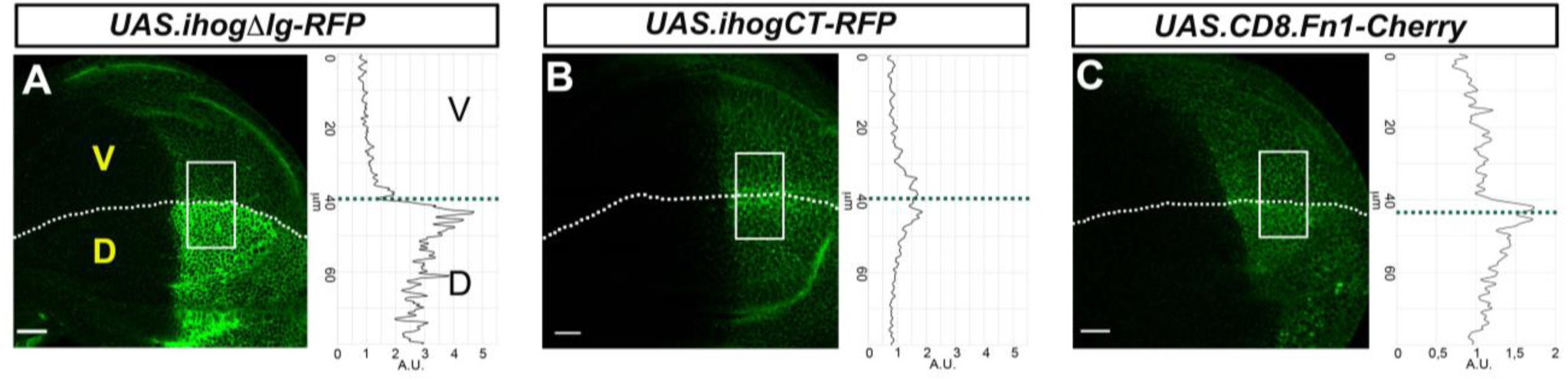
Role of Ihog domains for Hh retention. A-B) Wing disc*s* after 30 hours of induction at the restricted temperature of *apGal4TubGal80^ts^/+; UAS.ihogDIg-RFP/BacHhGFP* (A); *apGal4TubGal80^ts^/+; UAS.ihogCT-RFP/BacHhGFP* (B) and *apGal4TubGal80^ts^/+;UAS.CD8.Fn1-Cherry/BacHhGFP* (C). Plot profiles were performed over a ROI of size 80mm X 30mm. Note the build-up of Hh after the expression of IhogDIg-RFP (A) and the lack of accumulation of Hh after the expression of IhogDFN-RFP. Scale bar: 20 μm. Discs are oriented with dorsal part down (D), ventral up (V) and posterior right.

**Supplementary Figure 4.**
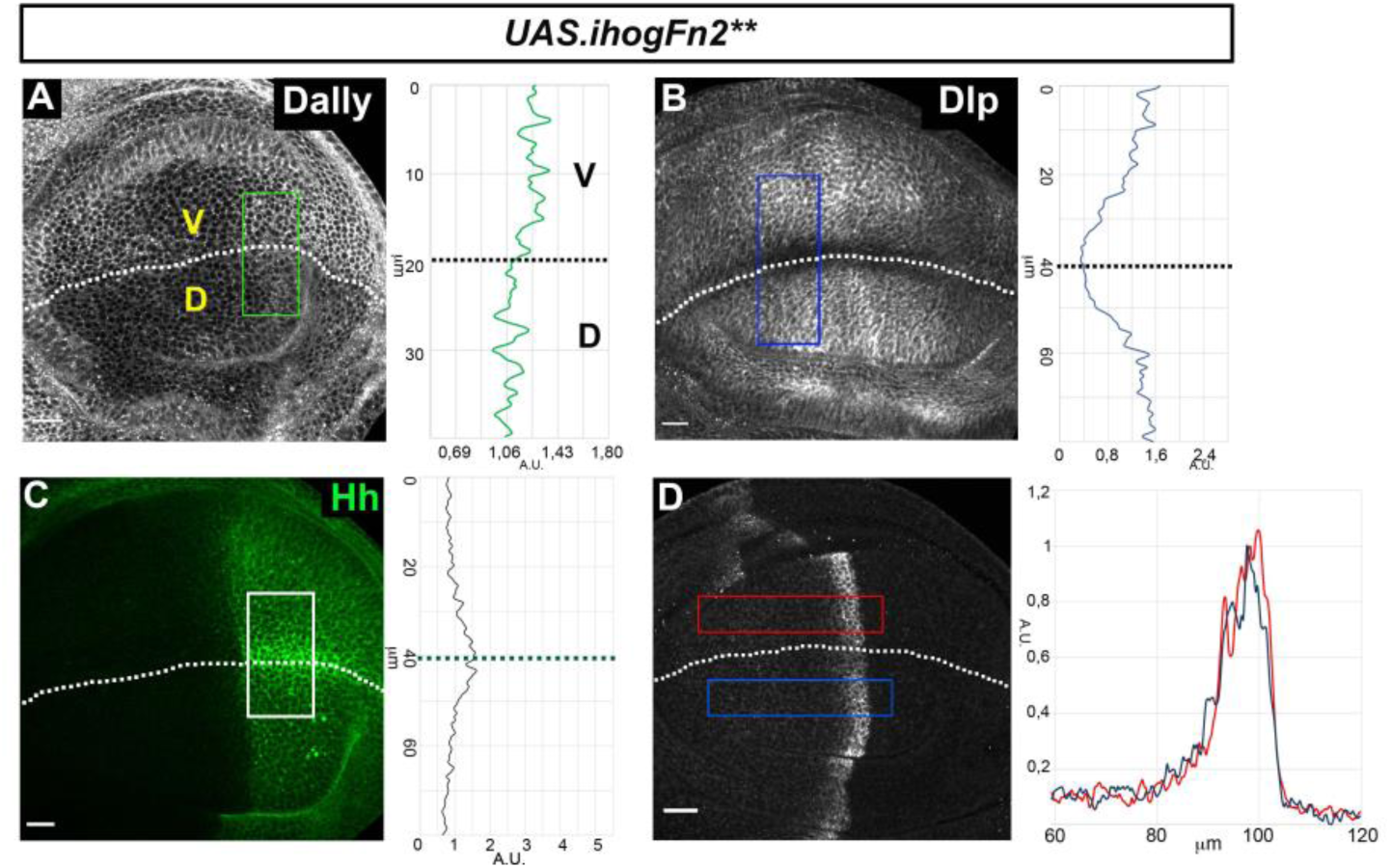
Effects of the ectopic expression of IhogFn2** (not tagged with RFP. A-D) Glypicans accumulation after 30 hours at the restricted temperature in *apGal4 tubGal80^ts^/ +; UAS.ihogFn2**/+* wing discs (A,B), Hh expression (BacHhGFP) in *apGal4 BacHhGFP/+; UAS.ihogFn2**-RFP/tubGal80^ts^* wing discs (C); Ptc expression in *LexAopGal80; apLexA ptcGal4/+; tubGal80^ts^/UAS.ihogFN2\*\** wing discs (D). Plot profiles where done over ROI of size 80μm X 30μm for Hh, 40μm X 20μm for Dally, 80μm X 20μm for Dlp and 120μm X 20μm for Ptc. Note that in all cases, its behavior is the same as that of IhogFn2**RFP.

**Supplementary Figure 5.**
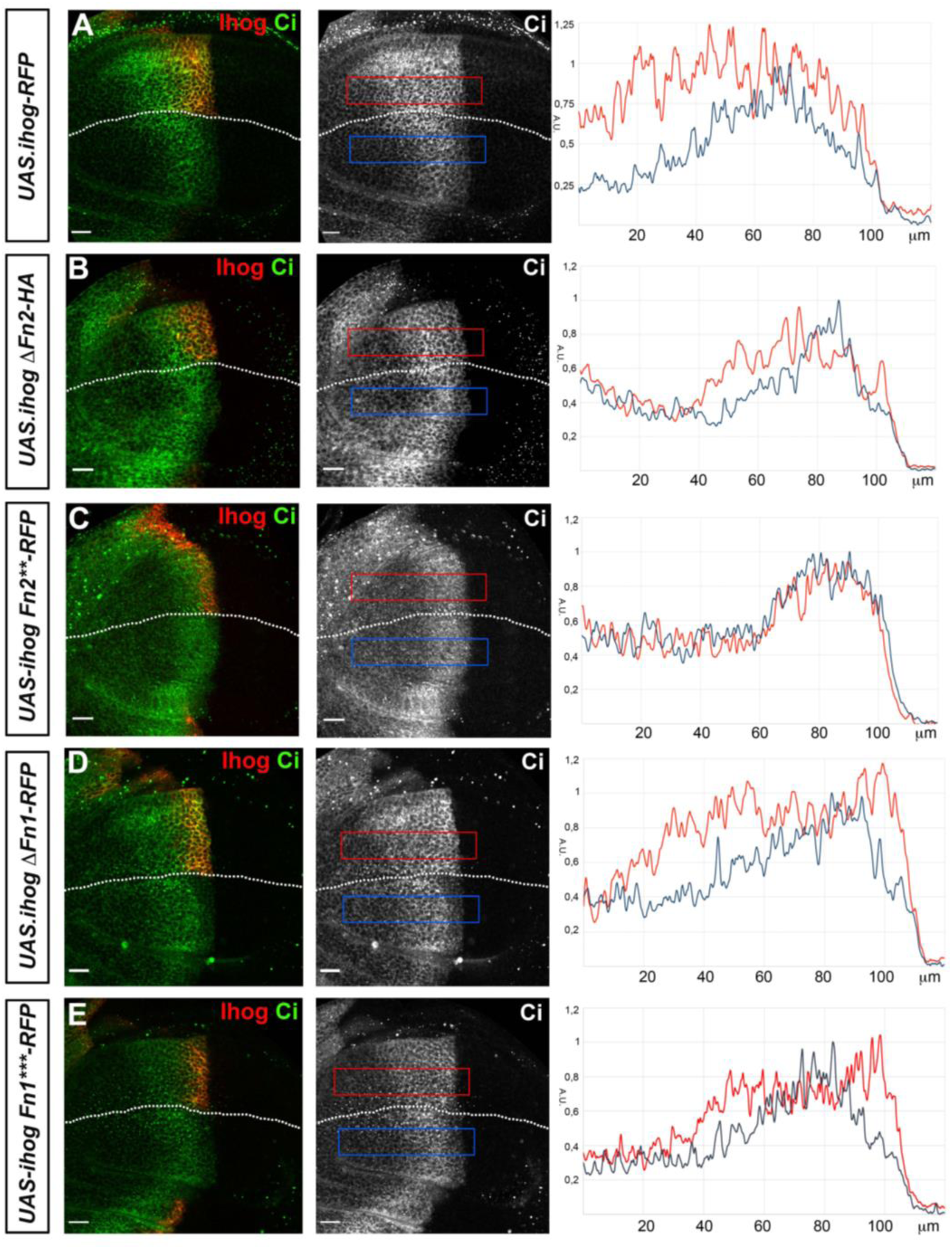
Role of the Ihog domains on Hh gradient formation (Ci). A-D) Wing disc Ci expression after induction of the different transgenes for 28 hours in *LexAopGal80; apLexA ptcGal4/+; tubGal80^ts^/+*: *UAS*.*ihog-RFP* (A); *UAS.ihogΔFn2-HA* (B); *UAS.ihogFn2**RFP* (C) and *UAS.ihogΔFn1-RFP* (D); and *UAS.ihogFn1***RFP* (E). Each image of the Ptc channel (red) incorporates plots of the fluorescence intensity in the ventral side of the wing disc. The dorsal side is used as internal control for Ci wild type expression (blue channel). Scale bar: 20 μm. Discs are oriented with dorsal part down and posterior right. Plot profiles were performed over a ROI of size 120μm X 20μm.

**Supplementary Figure 6.**
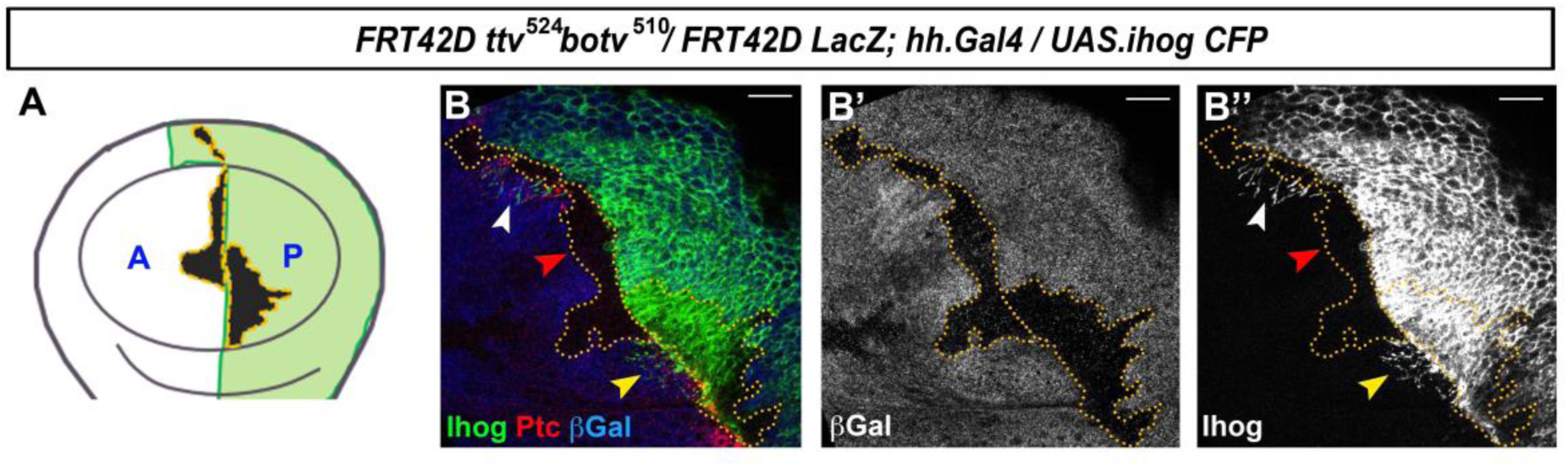
HSPG are needed for cytoneme stabilization in the adjacent cells. A-A’’) Behavior of cytonemes expressing Ihog in *ttv^524^ botv^510^* double mutant clones (labeled by the lack of β-gal, in blue) induced in a *hhGal4; UAS.ihogCFP* wing disc, with Ihog (in green) and Ptc (in red). Note that Ihog CFP expressing cytonemes can cross a small (white arrowhead) *ttv^524^ botv^510^* mutant clone induced the A compartment abutting the A/P border, but cannot cross when the clone is large (red arrowhead in A and A’’). Note also that cytonemes protruding from a *ttv^524^ botv^510^* mutant clone are stabilized (yellow arrow), indicating that HSPG are only needed for cytoneme stabilization in trans but not in cis.

**Supplementary Figure 7.**
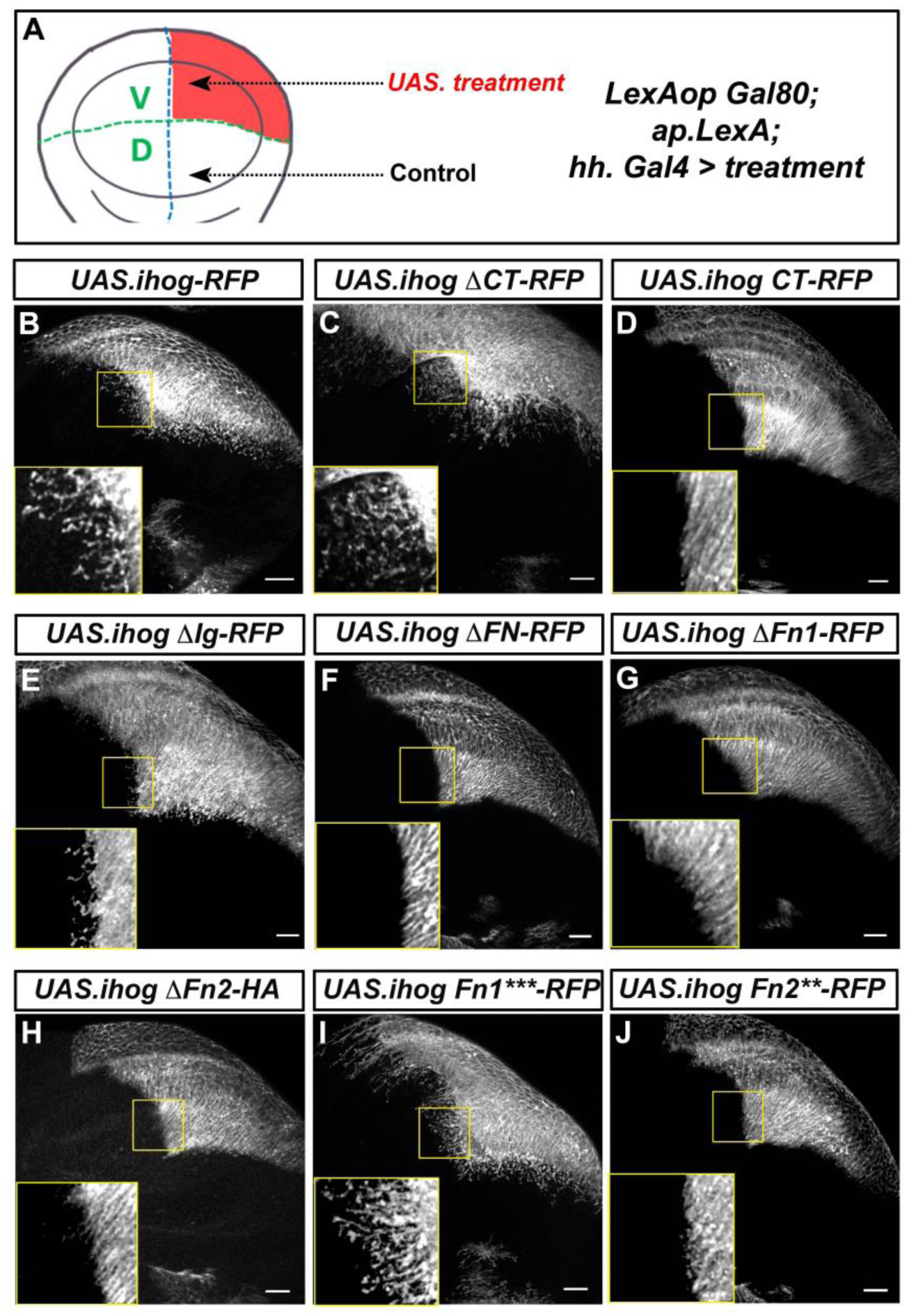
Ihog domains involved in cytoneme stabilization by ectopic expression in the wing disc. A) Scheme depicting the *LexAopGal80; apLexA; hhGal4* expression domain in the ventral part of the wing disc. B-J) Wing discs labeled cytonemes by expressing Ihog and Ihog mutant variants fused to RFP or HA after induction of the different transgenes for 28 hours in *LexAopGal80; apLexA/+; hhGal4TubGal80ts/+*: *UAS*.*ihog-RFP* (B); *UAS.ihogΔCT-RFP* (C); *UAS.ihogCT-RFP* (D); *UAS.ihogΔIg-RFP* (E); *UAS.ihogΔFN-RFP* (F)*;UAS.ihogΔFn1-RFP* (G); *UAS.ihogΔFn2-HA* (H); *UAS.ihogFn1***RFP* (I) and *UAS.ihogFn2**RFP* (J). An inset (yellow square) of 40×40 μm with amplified images are shown on the left side of each panel. Note long stabilized cytonemes by the expression of *UAS.ihog-RFP* (B), *UAS.ihogΔCT-RFP* (C) and *UAS.ihogFn1***RFP* (I). Note also that *UAS.ihogΔIg-RFP* (E) present very short cytonemes and that cytonemes are not visualized when the FNIII domains are deleted. Scale bar: 20 μm. Discs are oriented with dorsal part down and posterior right.

## Movie legends

**Movie 1**

**Cytoneme stabilization by ectopic Ihog require Glypican function in the neighboring cells.**

Abdominal histoblasts of pupae presenting *ttv^524^botv^510^* null mutant clones (abolishing glypicans function) in the A compartment, confronted to P compartment cytonemes marked with GMA-GFP (A) or with Ihog-CFP (B,C). *ttv^524^botv^510^* null clones are marked by the lack of RFP (lack of grey) and GMA-GFP or Ihog-CFP (in green). Cytonemes from the P compartment histoblasts labelled with GMA show the same dynamic behavior when traversing *ttv^524^botv^510^* null mutant clones than those traversing wild type territory (A). Cytonemes labelled with Ihog do not get stabilized when confronted to large *ttv^524^botv^510^* null mutant clones (B). Ihog-labelled cytonemes are stabilized and can cross small *ttv^524^botv^510^* mutant clones; it seems that the wild type neighboring cells can rescue the ectopic Ihog stability in cytonemes (C). Movies are 30 min long. Frames correspond to the projection of a Z-stack at 2 min intervals. The A compartment is on the left. Pupae were around 30 hours APF. Scale bar: 20 μm.

**Movie 2**

**Roles of Ihog domains on cytoneme stability**

Abdominal histoblasts expressing *UAS.GMA-GFP* (black) under the control of *Hh.Gal4 tubGal80^ts^* pupae (24 hours of expression) and also expressing *UAS.Ihog-RFP* (control) (A); *UAS.Ihog-ΔCT-RFP* (B); *UAS.IhogCT-RFP* (C); *UAS.Ihog-ΔIg-RFP* (D); *UAS.Ihog-ΔFN-RFP* (E); *UAS.Ihog-ΔFN1-RFP* (F) and *UAS.Ihog-ΔFN2-HA* (G). Note that cytonemes are stable in (A and B), while they are dynamic in (C-G). Movies are 30 min long. Frames correspond to the projection of a Z-stack at 2 min intervals. The A compartment is on the left. Pupae were around 30 hours APF. Scale bar: 10 μm.

**Movie 3**

**Different effect of Ihog-Hh and Ihog-Ptc interaction on cytoneme stability**

Abdominal histoblasts of pupae expressing *UAS.GMA-GFP* (in black) and *UAS.Ihog-RFP* (A); *UAS.Ihog-FN1***-RFP* (B); *UAS.IhogFN2**-RFP* (C); *UAS.IhogFN2\*\** (D) under the *Hh.Gal4 tubGal80^ts^* control (24 hours of induction). Note that cytonemes from cells expressing Ihog with alteration in its interaction with Hh (B) are as stable as those from the control expressing *Ihog-RFP* (A), while alteration of Ihog and Ptc interaction reverts cytoneme dynamics (C-D). Movies are 30 min long. Frames correspond to the projection of a Z-stack at 2 min intervals. The A compartment is on the left. Pupae were around 30 hours APF. Scale bar: 10 μm.

**Movie 4**

**Cytoneme dynamics of abdominal histoblasts expressing *UAS.boi-YFP***

Abdominal histoblasts of a *UAS.boi-YFP/+; Hh.Gal4 tubGal80^ts^* pupa expressing Boi-YFP (green) during 24 hours. (A) Z-stack from apical to basal, showing cytonemes in the most basal part. (B) 30 min movie of the same pupa showing dynamic cytonemes. Frames correspond to the projection of a Z-stack at 2 min intervals between frames. The A compartment is on the left. Pupa was around 30 hours APF. Scale bar: 10 μm.

**Movie 5**

**Cytoneme dynamics of histoblasts expressing both *UAS.boi-YFP* and *UAS.ihog-RFP***

Abdominal histoblasts of a *UAS.boi-YFP/UAS.ihog-RFP; Hh.Gal4 tubGal80^ts^* pupa. Boi-YFP (green) and Ihog-RFP (grey) were expressed during 24 hours before recording. (A) Z-stack from apical to basal showing basal cytonemes. (B) 30 min movie of the same pupa showing stabilized cytonemes. Frames correspond to the projection of a Z-stack at 2 min intervals. The A compartment is on the left. Pupa was around 30 hours APF. Scale bar: 10 μm.

**Movie 6**

**Cytoneme dynamics of histoblast expressing *UAS.ihog-RFP* and *UAS-RNAi-boi***

Abdominal histoblasts of a *UAS.Ihog-RFP/UAS.RNAi-Boi; Hh.Gal4 tubGal80^ts^*/+ pupa. Ihog-RFP (grey) and RNAi-Boi were expressed during 24 hours before recording. (A) Projection of a Z-stack from apical to basal show basal location of cytonemes. (B) A 30 min movie of the same pupa showing stabilized cytonemes. Frames correspond to the projection of a Z-stack at 2 min intervals. The A compartment is on the left. Pupa was around 30 hr APF. Scale bar: 10 μm.

**Movie 7**

**Different levels of Ihog and Boi in histoblasts affect cytoneme dynamics**

Abdominal histoblasts of pupae expressing *UAS.GMA-GFP* (black) under the control of *Hh.Gal4; tubGal80^ts^* and also expressing *UAS.Ihog-RFP* (after 24 hours of expression) (A). Note the stabilized cytonemes compared with the dynamic cytonemes of the control (expression of only GMA-GFP) (B). Abdominal histoblasts of a pupa continuously expressing *UAS.RNAi-Ihog UAS.RNAi-Boi* in the P compartment during development (C) showing cytonemes with similar dynamics than the control (B). Movies are 30 min long. Frames correspond to the projection of a Z-stack at 2 min intervals. The A compartment is on the left. Pupae were around 30 hours APF. Scale bar: 10 μm.

